# Phosphatidylserine clustering by membrane receptors triggers LC3-associated phagocytosis

**DOI:** 10.1101/2023.09.06.556449

**Authors:** Emilio Boada-Romero, Clifford S. Guy, Gustavo Palacios, Luigi Mari, Zhenrui Li, Douglas R. Green

**Affiliations:** Department of Immunology, St. Jude Children’s Research Hospital; Memphis, TN 38105, USA

## Abstract

LC3-associated phagocytosis (LAP) represents a non-canonical function of autophagy proteins in which ATG8 family proteins (LC3 and GABARAP proteins) are lipidated onto single-membrane phagosomes as particles are engulfed by phagocytic cells^1–4^. LAP plays roles in innate immunity^5^, inflammation and anti-cancer^6^ responses and is initiated upon phagocytosis of particles that stimulate Toll-like receptors (TLR), Fc-receptors, and upon engulfment of dying cells^6^. However, how this molecular route is initiated remains elusive. Here we report that receptors that engage LAP enrich phosphatidylserine (PS) in the phagosome membrane via membrane-proximal domains that are necessary and sufficient for LAP to proceed. Subsequently, PS recruits the Rubicon-containing PI3-kinase complex to initiate the enzymatic cascade leading to LAP. Manipulation of plasma membrane PS content, PS-binding by Rubicon, or the PS-clustering domains of receptors prevents LAP and phagosome maturation. We found that pharmacologic inhibition of PS clustering promotes the ability of dendritic cells to induce anti-cancer responses to engulfed tumor cells. Therefore, the initiation of LAP represents a novel mechanism of PS-mediated signal transduction upon ligation of surface receptors.

## Main Text

The process of phagocytosis, in which specialized cells such as macrophages engulf dead cells and/or pathogens, plays important roles in host defense, wound repair, and tissue homeostasis ^1,3^. This process depends on actin remodeling at the plasma membrane to form phagocytic “cups” that then seal to form phagosomes. These phagosomes mature through fusion with lysosomes to digest their cargo. LC3-associated phagocytosis (LAP) promotes phagosome maturation ^5^, important for innate defense against microbes ^7,8^, and its disruption promotes anti-cancer immunity ^6^. LAP utilizes components of the macro-autophagy (henceforth, autophagy) pathway to lipidate ATG8 family proteins onto the phagosome membrane, facilitating fusion with lysosomes ^1,3^. The process depends on the class III PI3-kinase VPS34 complex, and on the ligase machinery composed of ATG7, ATG3, and the complex of ATG16L and ATG5-12. Unlike autophagy, LAP functions independently of the ULK1/2 serine kinase complex and requires the protein Rubicon ^6^. LAP is initiated by engagement of cell surface Toll-like receptors (TLR) (including TLR1,2, TLR2,6, and TLR4), Fc-receptors, and receptors for dying cells ^1,3^, but the common signals shared by these receptors that function to initiate LAP are unknown.

During phagocytosis, the phospholipid phosphatidylserine (PS) concentrates in the cytosol-facing leaflet of the phagosome membrane ^9^ where it promotes binding of the kinase c-Src ^10,11^. We therefore considered that PS might also directly recruit the LAP machinery. We first asked whether signals that engage LAP preferentially enrich PS in the resultant phagosome. Beads coupled with the TLR2 ligand Pam3csk4 (Pam3, Pam3-beads) but not Biotin (control-beads) induce LAP ^5^, and therefore we purified phagosomes containing either cargo from immortalized bone marrow-derived macrophages (iBMDM) and performed lipidomic analysis. Principle component analysis revealed distinct profiles of lipids in phagosomes containing Pam3-coated beads (Pam3-phag) versus control, Biotin-coated beads (control-phag) (Fig. 1a, Extended Data Fig. 1a). Among the major lipid species, PS was enriched in Pam3-phag (Fig. 1b), an effect not accounted for by net increased PS (Extended Data Fig. 1b). To explore this in living cells, we expressed a fluorescent probe containing the C2 domain of Lactadherin (Venus-Lact-C2), which specifically binds PS ^9^, in RAW264.7 cells. Concordant with the lipidomic results, we observed recruitment of the probe to phagosomes containing Pam3-beads and IgG-coupled beads, (IgG-beads) but not control, BSA-coupled beads (Fig. 1c, Extended Data Fig. 1c). Similar results were obtained with another PS-binding probe based on the C2 domain of the clotting factor VIII (Venus-FVIII-C2) (Fig. 1d) ^12^. These results therefore suggest that the enrichment of PS in phagosomes depends on specific signals induced by the engulfed cargo.

**Figure 1:**
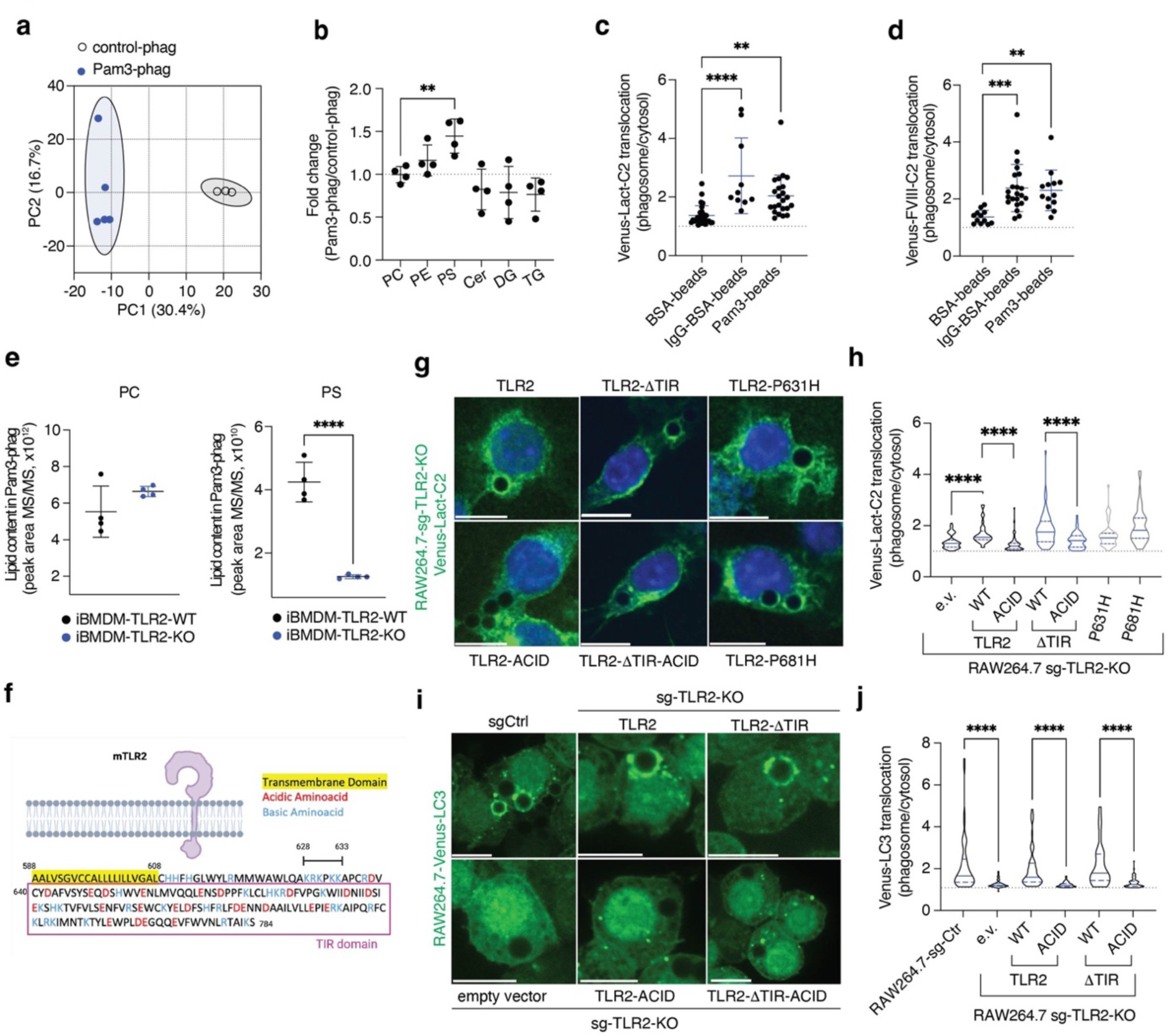
Phosphatidylserine species are enriched in the phagosome in a receptor-mediated manner independently of TIR signaling. (**a**) Principal component analysis of lipid content from phagosomes containing Pam3csk4-beads (Pam3-phag, blue) relative to phagosomes containing uncoupled-beads (control-phag, grey) isolated from immortalized bone marrow derived macrophages (iBMDM) in a representative experiment, n=4. (**b**) Fold change of MS/MS values of different lipid species in phagosomes containing Pam3-phag relative to control-phag isolated from iBMDM. Lipid species determined by lipidomics were aggregated per lipid class and each dot represents the cumulative value in n=4 independent experiments. PC, phosphatidylcholine; PE, phosphatidylethanolamine; PS, phosphatidylserine; Cer, Ceramide; DG, diacylglycerol; TG, triacylglycerol. (**c**, **d**) RAW264.7 cells stably expressing the PS-probes Venus-LACT-C2 or Venus-FVIII-C2 were fed BSA-beads, BSA-beads coupled with anti-BSA antibody (IgG-BSA-beads), or Pam3csk4-beads (Pam3-beads) for 30min and enrichment of PS-probe to phagosome membranes relative to cytosolic signal was determined by immunofluorescence. Each dot represents a phagosome (n>10) in a representative experiment, n=2. (**e**) Cumulative MS/MS peak area of PC and PS of Pam3-phag from wild-type (WT) and TLR2-KO iBMDM determined by lipidomic analysis. Replicates in a representative experiment, n=2. (**f**) Scheme of mouse TLR2 (Uniprot: Q9QUN7) depicted in a membrane. Transmembrane domain is highlighted in yellow; acidic and basic amino acid are colored in red and blue, respectively; and Toll-Interleukin receptor (TIR) domain is boxed in magenta. Numbers indicate amino acid position showcasing the basic patch (^628^KRKPKK^633^). (**g**-**j**) RAW264.7-sg-TLR2-KO cells stably expressing the PS-probe Venus-LACT-C2 (**g**, **h**) or Venus-LC3 (**i, j**) were transduced to express TLR2 full length or lacking the TIR domain (TLR2ΔTIR), in either wild-type (WT), K628E-R629D-K630E-K632E-K633E (ACID), P631H (corresponding to human SNP rs5743704), or P681H mutant TLR2. Transduction with empty vector (e.v.) served as a negative control. Cells were fed Pam3csk4-beads (30min, g, h; 1h, i, j) and Venus-LACT-C2 or Venus-LC3 translocation to phagosomes was determined by immunofluorescence. (g, i) Representative confocal images and (h, j) violin-plots of PS-probe enrichment at the phagosome membrane relative to cytosolic signal (n>20 phagosomes) in one representative experiment, n=2. \*\**P* < 0.01, \*\*\*\**P* < 0.001, \*\*\**P* < 0.0001 by two-sided Student’s t test.

Because Pam3 signals via TLR2 ^13^, we next asked if TLR2 is required for PS enrichment in response to this ligand. We generated TLR2-deficient iBMDM cells, exposed them to Pam3-beads, isolated phagosomes containing these beads, and performed lipidomic analysis. The presence or absence of TLR2 had no effect on cellular levels of PS (Extended Data Fig. 1d), but PS in the phagosomes of TLR2-deficient iBMDM was reduced as compared to phagosomes of WT iBMDM (Fig. 1e).

We noticed several positively charged residues (K, lysine; R, arginine) in the intracellular domain of TLR2 (TLR2-ID) near the transmembrane region and proximal to the TIR signaling domain (Fig. 1f) and speculated that these might interact with negatively charged PS. We therefore reconstituted TLR2-deficient RAW264.7 cells with wild-type TLR2 or TLR2 mutated at these five basic residues to acidic amino acids (^628^KRKPKK^633^ to ^628^EDEPEE^633^; TLR2^ACID^) (Extended Data Fig. 2a). After engulfment of Pam3-beads we assessed PS enrichment in the phagosome and found that the wild-type, but not the TLR2^ACID^ mutant, induced binding of the Venus-Lact-C2 probe to phagosomes (Fig. 1g,h), indicating that these basic residues (“basic patch”) are necessary for PS enrichment. We then deleted the entire TIR region with or without mutation of the basic patch and observed that even in the absence of the signaling domain, the basic patch of the TLR2-ID was necessary and sufficient for enrichment of PS, as assessed by binding of the PS probe (Fig. 1g,h). We further investigated this by making use of two previously reported mutations in TLR2. P631H corresponds to the human SNP rs5743704, that lies within the basic patch (^628^KRK**P**KK^633^) and is associated with reduced TLR2 signaling ^14,15^. P681H specifically reduces MyD88 signaling ^16^. Reconstitution of TLR2-deficient RAW264.7 with these TLR2 mutants (TLR2^P631H^, TLR2^P681H^; Extended Data Fig. 2a) did not affect PS enrichment (Fig. 1g,h). Therefore, TLR2-triggered PS enrichment in the phagosome appears to be independent of canonical MyD88 signaling, consistent with observations that the induction of LAP by TLR2 engagement is independent of MyD88 ^5^. LAP is not induced by Pam3-beads in TLR2-deficient cells ^5^, and we confirmed that while reconstitution of wild-type TLR2 restored recruitment of Venus-LC3B to phagosomes (Fig. 1i,j), TLR2^ACID^ failed to restore LAP in these cells, regardless of the presence or absence of the TIR signaling domain (Fig. 1i,j). Therefore, the requirements for PS enrichment and LAP induction appear to be the same.

We then asked if the intracellular domain of TLR2 (TLR2-ID, residues 609-784) interacts with PS. Recombinant TLR2-ID strongly bound immobilized PS and Cardiolipin compared with other lipids (Extended Data Fig. 2b,c). Furthermore, PS-coated beads precipitated the wild-type TLR2-ID (Fig. 2a), but mutation of the basic patch to alanine (Ala) or to acidic residues (D/E, Acid) abrogated PS binding (Fig. 2a). We then generated planar glass-supported lipid bilayers to be assayed by confocal microscopy. The lipid mixture resembled the lipid composition of the plasma membrane (see methods), and it included a fluorescently-labelled PS (TOP-FluorPS) and a nickel-containing lipid to favor proper orientation of N-terminal His-tagged recombinant proteins. After addition of His6X-TLR2-ID-FLAG to the lipid bilayer, we observed clustering of PS that was prevented by mutation of the basic patch or by addition of an anti-FLAG antibody (Fig. 2b,c). Taken together, it is likely that the clustering of PS by the basic patch of TLR2 contributes to the TLR2-dependent enrichment of PS in phagosomes containing Pam3-beads.

**Figure 2:**
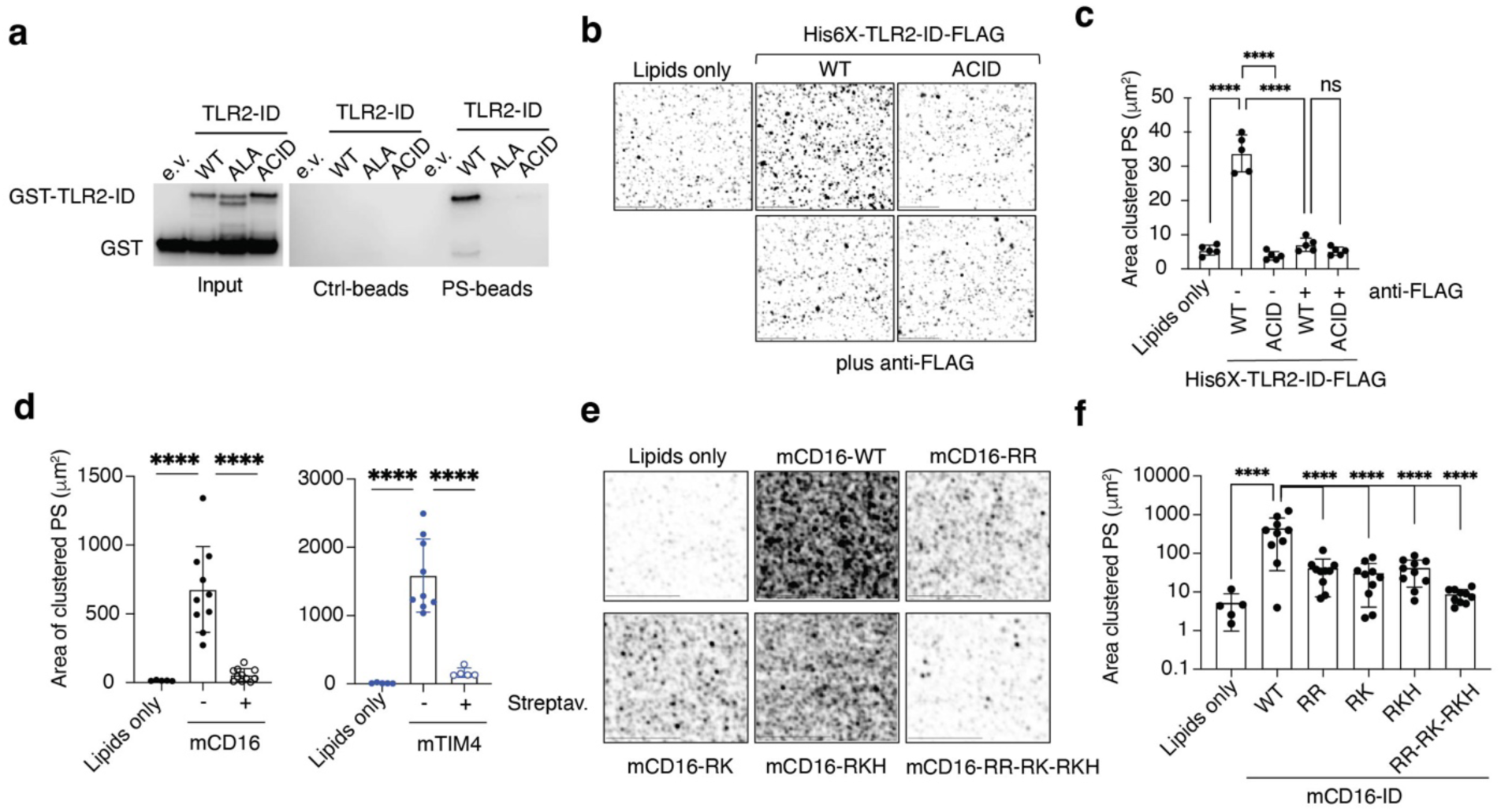
The cytosolic domains of receptors cluster phosphatidylserine. (**a**) PS-bead pull-down of TLR2 intracellular domain (TLR2-ID). Uncoupled-beads (Ctrl-beads), PS-beads, in combination with wild-type (WT), K628A-R629A-K630A-K632A-K633A (ALA), and K628E-R629D-K630E-K632E-K633E (ACID) versions were used as indicated, blots representative of n=4. (**b**-**f**) Glass-supported lipid bilayer resembling plasma membrane composition were incubated with TLR2-ID (b-c), cytosolic tails of mCD16 (d-f) or mTIM4 (d) as indicated, and top-Fluor-PS clustering was determined by immunofluorescence. (b, e) Representative images or (c, d, f) area of clustered PS quantified in different fields from representative experiments, n=2. (b-d) Anti-FLAG or streptavidin allows peptide manipulation and disrupts lipid clustering. \*\*\*\**P* < 0.001 by two-sided Student’s t test.

Next, we expanded our investigations to include receptors other than TLRs whose cargoes are also known to induce LAP, such as CD16, a component of the Fc receptor (FcR) that binds IgG ^17^, and Tim4 that binds dead and dying cells ^6^. IgG-coated beads induced enrichment of PS in the phagosome, as assessed by Venus-Lact-C2 recruitment (Fig. 1c,d, Extended Data Fig. 1c) and by lipidomics of isolated phagosomes (Extended Data Fig. 3a). Additionally, we generated N-terminal His-tagged intracellular regions of the CD16 subunit of FcR and of Tim4 coupled to biotin at the C-terminus (Extended Data Fig. 3b-d), and both peptides induced PS clustering in lipid bilayers that was disrupted by subsequent addition of streptavidin (Fig. 2d, Extended Data Fig. 3e). Like TLR2-ID, these peptides contain conserved basic patches despite a lack of sequence conservation between murine and human homologues (Extended Data Fig. 3c,f-g). We mutated these basic residues (lysine, K; arginine, R; and histidine, H), to acidic (aspartic acid, D; glutamic acid, E) and found that several of these (or all together) abrogated PS clustering (Fig. 1e,f). In addition, PS clustering in lipid bilayers with CD16-ID was dissipated with high salt (250mM NaCl vs. 50mM NaCl in conventional binding buffer) (Extended Data Fig. 3h,i). Therefore, receptors capable of engaging LAP have the property that basic residues in their intracellular regions cluster PS, most likely by electrostatic interactions.

Under homeostatic conditions, PS is predominantly localized to the inner leaflet of the plasma membrane by PS flippases ^18^, but this asymmetric distribution can be disrupted by phospholipid scramblases ^19,20^. To ask whether the observed enrichment of PS in the phagosome membrane is required for LAP, we took advantage of the ability of calcium ionophores, such as ionomycin, to induce phospholipid scrambling ^19^ and then “locked” PS on the outer leaflet with a PS-specific antibody (Fig. 3a,b; Extended Data Fig. 3a). This technique effectively diminished phagosome PS levels, as detected by a PS probe, upon feeding RAW264.7 cells with yeast particles (zymosan, another LAP inducer ^5^; Extended Data Fig. 3b,c). RAW264.7 cells expressing Venus-LC3B (RAW264.7-Venus-LC3) were similarly treated with ionomycin together with control or anti-PS antibody and then fed zymosan. Recruitment of Venus-LC3B to phagosomes was detected by confocal microscopy or by FACS wherein the Venus signal is retained following cellular permeabilization with digitonin ^21^. These assays revealed that while zymosan induced recruitment and retention of LC3 to phagosomes (LAP) in ionomycin-treated cells under control conditions (no antibody or IgG control), ionomycin with anti-PS antibodies prevented LAP induced by zymosan (Fig. 2c,d).

**Figure 3.**
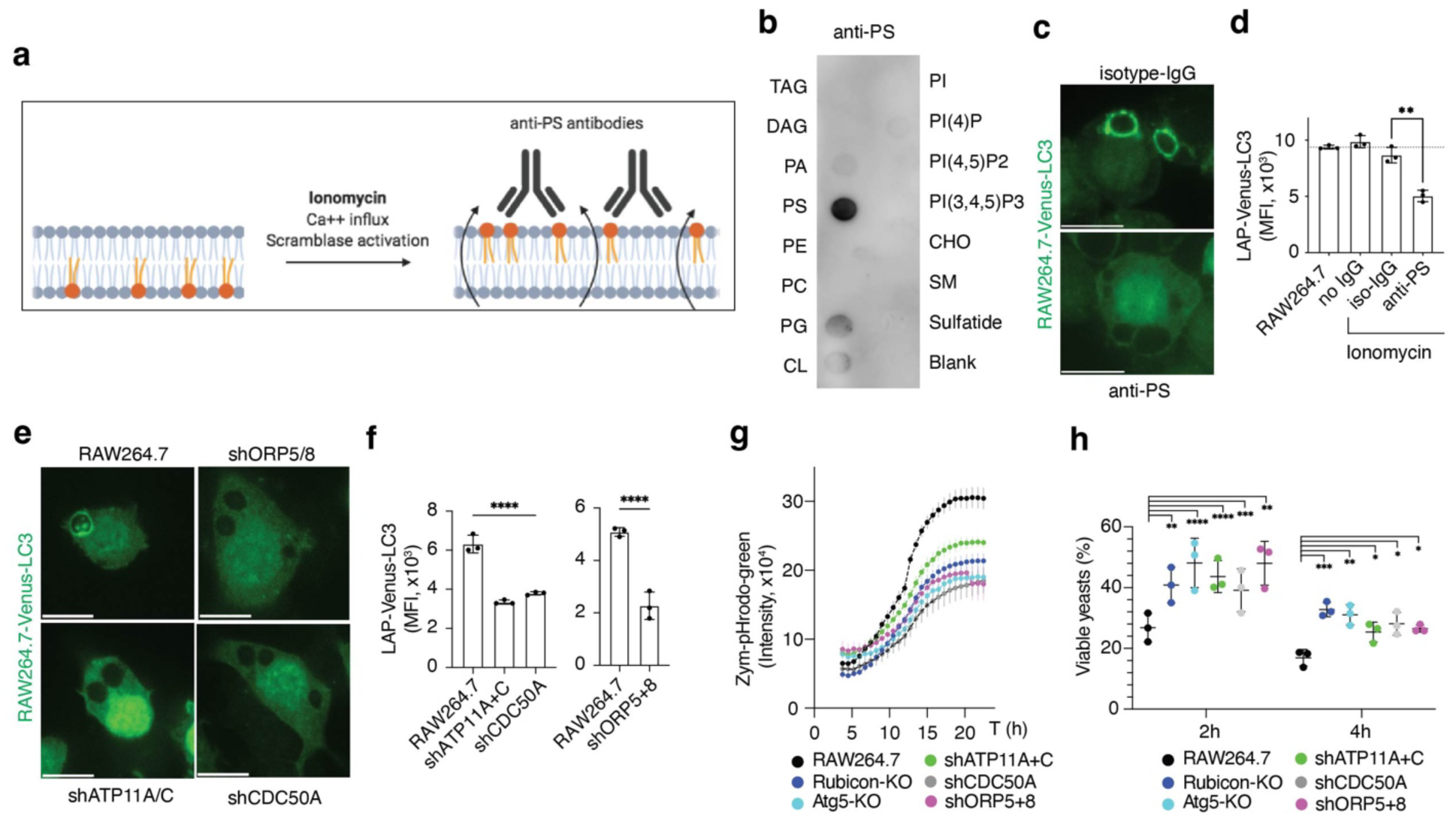
Reduced phosphatidylserine levels impair LC3-associated phagocytosis. (**a**) Scheme showing the phosphatidylserine (PS) trapping strategy using anti-PS antibody. At the steady-state, PS is confined to the cytosolic leaflet at the plasma membrane. The calcium ionophore ionomycin induces a flux of calcium into the cells that in turns activates calcium-dependent lipid scramblases that expose PS at the cell surface. Specific anti-PS antibodies lock the PS at the outer leaflet and preclude its localization in the cytosolic face. (**b**). Lipid strip showing the binding specificity of anti-PS antibody. PS: phosphatidylserine; PA: phosphatidyl acid; PG: phosphatidylglycerol; PC: phosphatidylcholine; PE: phosphatidylethanolamine; DAG: diacyl-glycerol; TAG: triacyl-glycerol; CL: cardiolipin; SM: sphingomyelin; CHO: cholesterol; PI: phosphatidylinositol. (**c, d**) RAW264.7 cells stably expressing Venus-LC3 cells (RAW264.7-Venus-LC3) were treated with ionomycin (10mM, 30min) combined with anti-phosphatidylserine (PS) antibody (1:50) or isotype control (anti-FLAG, 1:50) and fed Zymosan (1h). (**e, f**) RAW264.7-Venus-LC3 cells silenced for ATP11A and ATP11C, CDC50A, or ORP5 and ORP8 using a stably transduced short hairpin (sh) RNA (shATP11A/C, shCDC50A, and shORP5/8,) and fed Zymosan (1h). (c, e) Representative confocal images and (d, f) Venus-LC3 levels in Zymosan-TexasRed^+^ cells after digitonin treatment to assess retained Venus-LC3. (**g**) Phagosome acidification over time upon feeding with acid-sensitive probe Zymosan-pHRodo-Green in the indicated cell lines. (**h**) Yeast killing capacity after 2h and 4h of yeast engulfment. Values normalized to yeast recovered after 1h of engulfment per cell line (time 0h). RAW264.7-Atg5-KO and Rubicon-KO cells display LAP deficiency. Data are means ± SD of three (d, f, h) or eight (g) biological replicates. Each representative of 3 independent experiments. \**P* < 0.05, \*\**P* < 0.01 \*\*\**P* < 0.001 \*\*\*\**P* < 0.001 by two-sided Student’s t test.

As a second approach to reduce PS enrichment, we silenced the PS flippases (Extended Data Fig. 5a) ATP11A and ATP11C or their requisite chaperone, CDC50A ^22^ that are abundant in mouse macrophages (Extended Data Fig. 5b). Effective silencing (Extended Data Fig. 5c,d) of these proteins impaired the enrichment of PS on the phagosome membrane (Extended Data Fig. 5h,i) and prevented the retention of Venus-LC3B following phagocytosis of zymosan (Fig. 3e,f).

PS is synthesized in the endoplasmic reticulum and transported to the plasma membrane by the Oxysterol-binding related proteins 5 and 8 (ORP5 and ORP8), which exchange PS for phosphoinositide-4P (PI4P), generated from PI by PI4KIIIa in the plasma membrane ^23^ (Extended Data Fig. 5e). Silencing of ORP5 and ORP8 in RAW264.7 cells (Extended Data Fig. 5f,g) reduced PS levels at the phagosome membrane (Extended Data Fig. 5h,i) and prevented the retention of Venus-LC3B upon engulfment of zymosan (Fig. 3f,g). Altogether, our results support the idea that enrichment of PS in the phagosome membrane promotes LAP.

LAP promotes phagosome maturation ^5,7,8^, and we therefore interrogated the downstream acidification of phagosomes using zymosan labeled with the pH-sensitive dye pHrodo ^24^. Zymosan-containing phagosomes acidified in wild-type RAW264.7 cells, but this was reduced in LAP-deficient RAW264.7 cells lacking Rubicon or ATG5 (Fig. 3g). Testing the role of PS enrichment in this process, we found that cells in which ORP5 and ORP8, ATP11A and ATP11C, or CDC50A were silenced showed similarly delayed phagosome acidification (Fig. 3g). Ablation of ATG5 or Rubicon also delayed the killing of yeast (Fig. 3h), as did silencing of ORP5 and ORP8, ATP11A and ATP11C, or CDC50A (Fig. 3h), consistent with the role of LAP in phagosome maturation ^7,8^.

We next pharmacologically reduced PS plasma levels, employing a PI4KIIIa inhibitor, GSK-A1^23,25^, and an inhibitor of PS levels at the plasma membrane, Fendiline ^26^. Consistent with other reports, GSK-A1 reduced cellular levels of PS ^23^ (Extended Data Fig. 6a), and Fendiline reduced ceramide levels and reduced plasma membrane PS ^26^. Treatment of RAW264.7 cells with either inhibitor reduced PS enrichment in zymosan-containing phagosomes (Extended Data Fig. 6b,c), and LC3 translocation to phagosomes (Fig. 4a,b). Phagosome acidification was similarly inhibited to the same extent as in Rubicon-deficient cells (Fig. 4c). In contrast, these inhibitors did not affect canonical autophagy (Extended Data Fig. 6d). Additionally, these inhibitors reduced BMDM killing of yeast (Extended Data Fig. 7a,b), consistent with the idea that PS enrichment in the phagosome membrane supports lipidation of LC3 proteins to promote maturation and acidification.

**Figure 4:**
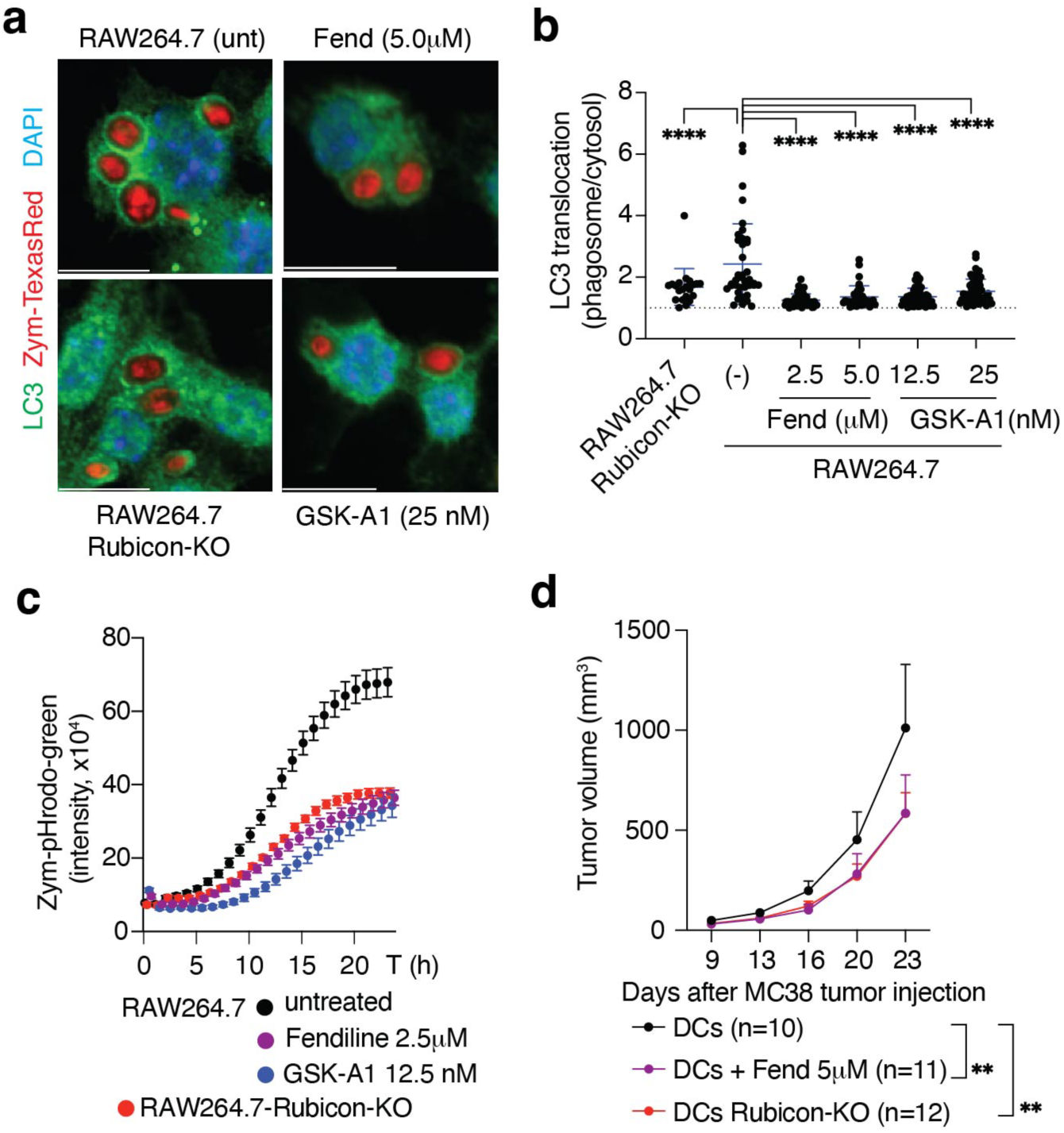
Chemical inhibition of PS metabolism serve as LAP inhibitor. (**a, b**) RAW264.7 were pretreated as indicated for 16h and fed Zymosan-TexasRed particles. Representative confocal images (a) and cumulative data (b) of endogenous LC3 enrichment at the phagosome membrane relative to cytosolic signal in n>10 phagosomes. (**c**) Phagosome acidification over time in RAW264.7 cells pretreated with inhibitors (16h) upon feeding Zymosan-pHRodo-Green. Data are means ± SD eight biological replicates, representative of 3 independent experiments. (**d**) C57BL/6J mice were implanted subcutaneously with 10^5^ MC38 cells. CD8^+^ DCs were treated *ex vivo*, or not with Fendiline (5μM, 20h) as indicated, and co-cultured with killed MC38 cells for 3h. At days 7 and 14 post-implantation mice were injected intradermally with 10^6^ of these CD8^+^ DCs and tumor growth assessed. Data are mean + SEM of n>10 mice in 2 independent experiments. \*\**P* < 0.01 \*\*\*\**P* < 0.001 by two-sided Student’s t test (b) or ANOVA test (d).

The ability to engage LAP is associated with inhibition of anti-tumor immune responses, subcutaneously implanted tumor cells generated smaller tumors in LAP-deficient mice compared to LAP-proficient mice, a phenomenon dependent on adaptive immunity but stemming from the myeloid compartment ^6^. To test the effect of LAP-specific inhibitors in an in vivo setting we pre-treated conventional dendritic cells (DCs) with Fendiline and primed them with dead MC38 cells *ex vivo*, and then transferred them to MC38 tumor-bearing mice. DCs that were treated with the PS inhibitor induced a delay in tumor growth comparable to the transfer of Rubicon-deficient DCs, as compared to untreated, primed WT DCs (Fig. 4d), supporting the use of LAP inhibitors as anti-tumor promoting agents.

Like autophagy, LAP requires the class III PI3-kinase VPS34 ^6^ and its product, phosphatidylinositol 3-phosphate (PI3P). LAP also requires Rubicon, but while Rubicon is not generally required for VPS34 activity or LC3-lipidation during canonical autophagy ^27,28^ (Extended Data Fig. 8a), VPS34 activity upon LAP requires Rubicon (Extended Data Fig. 8a,b). PI3P can be detected with probes based on the PI3P-binding PX domain of p40phox ^29,30^ (Extended Data Fig. 8c,d). Using either recombinantly produced (Extended Data Fig. 8e,f) or genetically encoded fluorescent probes (Extended Data Fig. 8g,h), we determined that the binding of PX-p40phox to zymosan-containing phagosomes was absent in Rubicon-deficient cells.

Components of the PI3KC3 complex, including VPS34, VPS15, and Beclin-1, co-precipitate with Rubicon ^27,28^ (Extended Data Fig. 8i). PI3KC3 complex translocation as well as LC3-lipidation were reduced or absent in phagosomes from Rubicon-deficient cells (Extended Data Fig. 8j). Rubicon-deficient cells reconstituted with full length FLAG-Rubicon co-precipitated PI3KC3 complex components (Extended Data Fig. 9a), that were enriched at the phagosome membrane (Extended Data Fig. 9b). Immunoprecipitation of FLAG-Rubicon from Pam3-phag lysates revealed co-precipitation of the PI3KC3 complex (Extended Data Fig. 9c) and VPS34 activity, detected by PI3P generation *in vitro* (Extended Data Fig. 9d). Rubicon binds the PI3KC3 complex via its coiled-coil domain (CCD) ^27,28,31^, and deletion of this domain from FLAG-Rubicon (RubiconΔCCD) prevented co-precipitation of the PI3KC3 complex (Extended Data Fig. 9a), translocation of PI3KC3 partners to phagosomes, and VPS34 activity on phagosomes (Extended Data Fig. 9b-d). These results support the idea that Rubicon is required for recruitment and PI3KC3 complex activity at phagosome membranes.

Despite reduced interaction with PIK3C3, RubiconΔCCD translocated to phagosomes (Extended Data Fig. 9a,b,e,f), suggesting that Rubicon recruitment and PIK3C3 interaction are independent events. We found that recruitment of Rubicon to phagosomes was limited when PS was reduced at the phagosomes (Fig. 5a,b), in the absence of TLR2, or in the presence of TLR2^ACID^ or TLR2ι1TIR^ACID^ (both unable to interact with PS); however, it was independent of canonical TLR2 signaling mediated by the TIR domain (Fig. 5c, Extended Data Fig. 10a). In contrast, ablation of Rubicon did not affect the recruitment of Venus-Lact-C2 to phagosomes (Extended Data Fig. 10b,c). Therefore, Rubicon recruitment to phagosomes is dependent upon PS enrichment and occurs downstream of this event.

**Figure 5:**
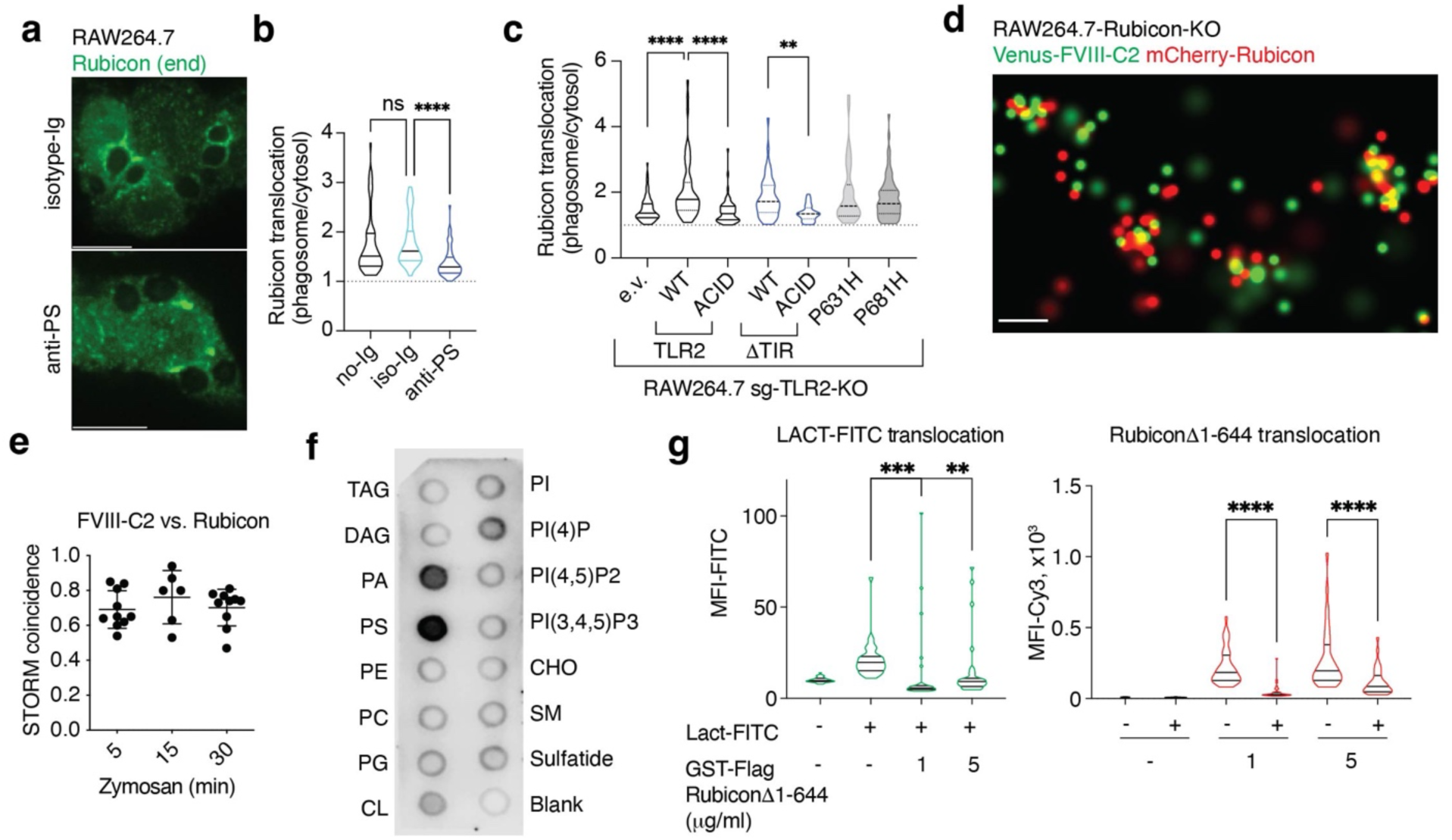
Rubicon binds to phosphatidylserine in the phagosome membrane. (**a, b**) RAW264.7 cells were treated with ionomycin (10μM, 30min) combined with anti-phosphatidylserine (PS) antibody (1:50) or isotype control (anti-FLAG, 1:50) and fed Zymosan (30min). Representative confocal images (a) and violin-plots (b) of endogenous Rubicon enrichment at the phagosome membrane relative to cytosolic signal (n>40 phagosomes). (**c**) RAW264.7-sg-TLR2-KO cells were transduced to express full length TLR2 or TLR2 lacking the TIR domain (TLR2τιTIR), in either wild-type (WT), K628E-R629D-K630E-K632E-K633E (ACID), P631H (corresponding to human SNP rs5743704), or P681H mutant TLR2. Transduction with empty vector (e.v.) serves as a negative control. Cells were fed Pam3csk4-beads (30min) and Rubicon translocation was determined by immunofluorescence. Violin-plots showing the enrichment of Rubicon at the phagosome membrane relative to the cytosolic level (n>20 phagosomes). (**d, e**) RAW264.7-Rubicon-KO cells stably expressing the PS-probe Venus-FVIII-C2 and mCherry-Rubicon were fed Zymosan and analyzed by stochastic optical reconstruction microscopy (STORM). (d) Super-resolution image showing a representative membrane portion of a Zymosan-containing phagosome and (e) statistical index in super-resolution images assessing the proximity of PS-probe and Rubicon at phagosome membranes over time. Data are means ± SD of >6 biological replicates (phagosomes) in one representative experiment. (**f**) Lipid strip showing the lipid-binding specificity of full-length Rubicon recombinantly produced in insect cells, n=2. (**g**) *In vitro* binding competition assays of proteins in Pam3csk4-beads containing phagosomes isolated from RAW264.7-Rubicon-KO cells. Phagosomes were incubated with Lactadherin-FITC (Lact-FITC) and/or GST-Rubicon D1-644 as indicated. After anti-GST-Cy3 staining, phagosomes were quantitively analyzed by microscopy for green (Lactadherin binding) or red signal (GST-Rubicon Δ1-644 binding). Violin plots depict n>30 phagosomes. ** *P* <0.01, *** *P* <0.001, **** *P* <0.0001 by (b, g) two-sided Student’s t test or (c) ANOVA test (pairwise comparations Fisher’s LSD).

To observe the interplay between Rubicon and PS in living cells, we reconstituted Rubicon-deficient RAW264.7 cells with mCherry-Rubicon. These cells also expressed Venus-FVIII-C2 or Venus-Lact-C2 to visualize PS enrichment on phagosomes. Using either probe, we observed co-recruitment of Rubicon and PS probes to phagosomes upon engulfment of Pam3-beads or BSA-beads with anti-BSA antibody (i.e.: Ig-coated Beads), but not with BSA-beads alone (Extended Data Fig. 11a-d). Super-resolution microscopy revealed a strong co-association of Rubicon and PS at the phagosome membrane (Fig. 5d,e; Extended Data Fig. 11e,f), and recombinant Rubicon preferentially bound to both phosphatidic acid (PA)- and PS-immobilized lipids (Fig. 5f, Extended Data Fig. 12a). To determine the molecular basis of Rubicon translocation, we generated two fragments of Rubicon, an N-terminal fragment to residue 644 and a C-terminal fragment beginning at 645. While only the C-terminal fragment was recruited to zymosan-containing phagosomes (Extended Data Fig. 12b), only the N-terminal fragment (containing the CCD), bound the PI3KC3 complex (Extended Data Fig. 12c). Unlike full-length Rubicon, neither fragment showed VPS34 activity in phagosomes (Extended Data Fig. 12d). Therefore, it is likely that the C-terminal region mediates binding to phagosomes whereas the N-terminal region recruits the PI3KC3 complex. To test if the C-terminal region binds PS, we generated the recombinant C-terminal fragment and examined its binding to isolated phagosomes in the presence or absence of recombinant Lactadherin, which binds specifically to PS ^32^. We found that the C-terminal fragment of Rubicon and Lactadherin competed for binding to phagosomes (Fig. 5g), further supporting the idea that the C-terminal region of Rubicon binds PS on phagosomes.

The C-terminal region of Rubicon contains a FYVE-like domain ^33^, which in other proteins binds PI3P ^34^. We found that mutating basic amino acids in this domain (718-721; KRLR to EELE; Rubicon^MUT^; Fig. 6a) dramatically reduced the co-precipitation of lipid species with Rubicon following zymosan engulfment (Extended Data Fig. 13a). Despite similar PI3KC3 complex binding (Fig. 6b), Rubicon^MUT^ was not recruited to phagosomes (Fig. 6c,d) and did not promote generation of PI3P at the phagosome membrane (Fig. 6e; Extended Data Fig. 13b). Furthermore, while Rubicon restored efficient LAP (Fig 6f; Extended Data Fig. S13c) and phagosome maturation to Rubicon-deficient cells, neither Rubicon^MUT^ nor the C-terminal fragment of Rubicon supported phagosome maturation (Fig. 6g; Extended Data Fig. 13d). Our results suggest that Rubicon interacts with the PI3KC3 complex and recruits this activity to PS-enriched phagosomes to promote LAP and phagosome maturation.

**Figure 6:**
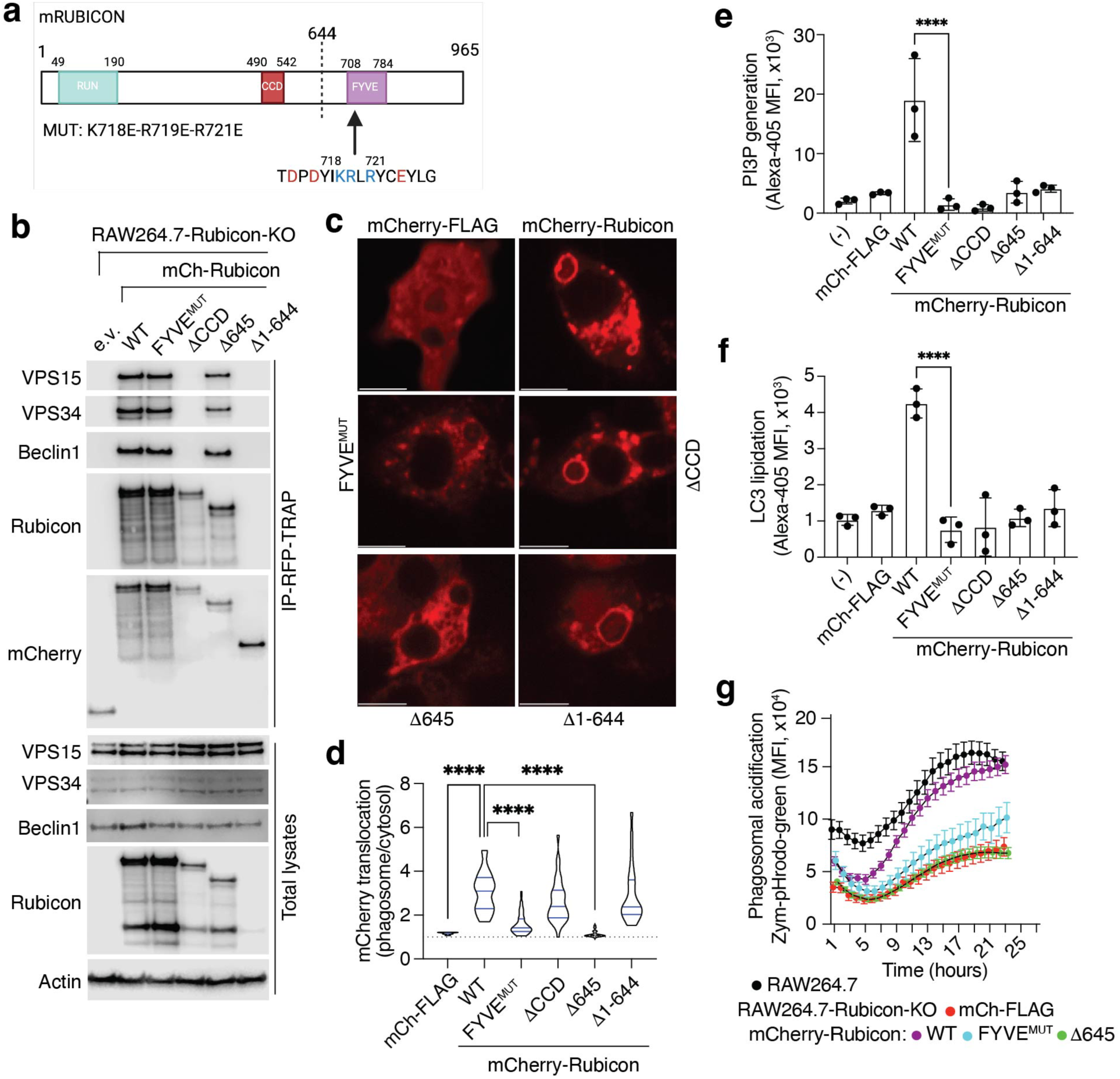
Binding of Rubicon to phosphatidylserine via FYVE domain is essential for LAP. (**a**) Scheme of mouse Rubicon protein (Uniprot: Q80U62). RUN: ṞPIP8, U̱ NC-14 and ṈESCA domain; CCD: coiled-coil domain; FYVE: Fab-1, YGL023, Vps27, and EEA1 domain. Numbers indicate amino acid position and highlighted are amino acids in the lipid binding core within the FYVE domain. (**b**) RAW264.7-Rubicon-KO cells stably expressing mCherry-tagged Rubicon (mCh-Rubicon), WT or FYVE^MUT^, 1′CCD, 1′645, or 1′1-644, were lysed, subjected to RFP-TRAP-immunoprecipitation (IP) and immunoblotted. Total lysates serve as input for IP. FYVE^MUT^: K718E-R719E-R721E and 1′CCD (1′Coiled-coil domain, lacks Beclin1-interacting region, amino acids 490-542). (**c**, **d)** RAW264.7-Rubicon-KO cells stably expressing different versions of mCherry-Rubicon were fed Zymosan (30 minutes). Representative confocal images (c) and violin-plots (d) showing enrichment of mCherry signal at the phagosome membrane relative to the cytosol (n>25 phagosomes). Representative of 2 independent experiments. (**e**, **f**) Cytometric analysis of PI3P generation or LC3-lipidation in RAW264.7-Rubicon-KO cells expressing different versions of mCherry-Rubicon. PI3P was detected with recombinant GST-p40-phox-PX (d) or immobilized LC3 levels (e) in Zymosan-FITC^+^ cells after digitonin treatment. (**g**) Phagosome acidification over time in RAW264.7 or RAW264.7-Rubicon-KO cells expressing different versions of mCherry-Rubicon upon feeding with the acid-sensitive probe Zymosan-pHRodo-Green. Data are means ± SD of 3 (e, f) or 8 (g) biological replicates. Each representative of 2 (g) or 3 (e, f) experiments. mCherry-FLAG is used as negative control, (-) indicates untransduced RAW264.7-Rubicon-KO cells (e, f). \*\*\*\**P* < 0.0001 by two-sided Student’s t test.

Based on our findings, we propose the following model for the initiation of LAP (Extended Data Fig. 14a). Receptors containing the basic patch, when ligated, enrich PS in the phagosome as it forms. The Rubicon-containing VPS34-complex is then recruited to phagosomes via the binding of Rubicon to PS. This generates PI3P on the phagosome membrane, which, in turn, recruits the E3-ligase complex, ATG16L-5-12 ^35,36^ to lipidate ATG8 family proteins on the single phagosome membrane. The phagosome is now decorated with these ATG8 proteins, facilitating fusion with lysosomes to digest the cargo ^5^.

During canonical autophagy, ATG8 is exclusively ligated to phosphatidylethanolamine (PE). However, during noncanonical conjugation of ATG8 to single membranes (CASM), ATG8 proteins are ligated to both PE and PS ^37^. It is possible that this ligation to PS is facilitated by the enrichment of PS in phagosomes during the CASM process of LAP. Another CASM process is LC3-associated endocytosis (LANDO) ^4,38^, in which ATG8 proteins might also conjugate to both PE and PS ^37^. Some endosomes have also been found to be enriched for PS ^39^, and it is therefore possible that a similar mechanism to what we have described for LAP exists for such PS enrichment, Rubicon binding, and ATG8 lipidation during LANDO. Unlike LAP, LANDO does not appear to promote endosome fusion to lysosomes, but instead promotes recycling of some receptors from the endosome to the plasma membrane ^4,38^. Interestingly, we found that cells in which ORP5 and ORP8, ATP11A and ATP11C, or CDC50A were silenced showed similarly delayed recycling of TREM2 receptor in RAW264.7 cells as ATG5 deficiency (Extended Data Fig. 14b,c). TREM2 has a potential basic patch in its cytosolic tail (Extended Data Fig. 14d), suggesting that a similar mechanism may take place during LANDO.

LAP and LANDO have important roles in innate immunity, anti-cancer immunity, and neurodegenerative disease ^4,38^ and are likely to have additional roles in other physiologic and pathologic settings. While protein-lipid interactions are well known to play roles in different signal transduction pathways, we suggest that the interactions we have described here, where a signal is transduced from PS-clustering receptors (e.g. TLR2, CD16, TIM4) to downstream PS-binding proteins (e.g. Rubicon in the PI3KC3 complex), represents a novel form of signal transduction. It is notable, though, that PS-enrichment on phagosomes engages c-Src ^10,11^, and we can therefore envision that the activation of c-Src upon engagement of TLR2 may depend on the ability of the latter to bind and cluster PS. Thus, we expect there will be additional pathways initiated in a manner similar to the one described here.

## Methods

### Mice

Rubicon-KO mice were previously described ^40^, TLR2-KO mice were obtained from the Jackson Laboratories (Cat. No. 004650). Rubicon and TLR2 mouse lines were backcrossed to transgenic reporter mice expressing GFP-LC3 ^41^ (generous gift from Dr. Noboru Mizushima) on the C57BL/6J background. Female recipient C57BL/6J mice for dendritic cell transfers were purchased from Jackson Laboratories (Cat. No. 000664). Age- and sex-matched littermates were used as controls. Mice were bred and housed in pathogen-free facilities, in a 12-hour light/dark cycle in ventilated cages, with chow and water supply ad libitum at the Animal Resources Center in St. Jude Children’s Research Hospital. Procedures were approved by the Institutional Animal Care and Use Committee at St. Jude and in compliance with all relevant ethical guidelines.

### Cell lines and culture conditions

Cancer cell lines HEK293T (Cat. No.: CRL-3216), RAW264.7 (Cat No. TIB-71), and L-929 (Cat. No.: CCL-1) were purchased from ATCC. MC38 (mouse colon carcinoma) cells were a gift from Dr. Hongbo Chi (St. Jude Children’s Research Hospital, Memphis, TN, USA), B16F10 (mouse melanoma) cells secreting Fms-related tyrosine kinase 3 ligand (FLT3L; B16F10-FLT3L cells) were a gift from Dr. Steven S. Porcelli (Albert Einstein College of Medicine, New York, NY, USA) ^42^. RAW264.7-Rubicon-KO and RAW264.7-ATG5-KO cells were generated by CRISPR-Cas9 as previously described ^2,6^; these cells were used to benchmark LAP-deficiency in most of the experiments. RAW264.7-TLR2-KO cells were generated by CRISPR-Cas9 using the LentiCRISPR-v2 system (kind gift from Dr. Brett Stringer, Addgene#98290) targeting an early PAM site in the Tlr2 coding exon. After lentiviral transduction, cells were selected by repetitive cell sorting using a validated monoclonal antibody (Clone T2.5, Biolegend 121810) until a homogenous polyclonal negative population was obtained. Lack of TLR2 expression in the negative population was confirmed by western blot using a different antibody (Cell Signaling Technology-CST, 13744). Re-expression of mTLR2 or its mutants was achieved by transduction of negative cells with retrovirus carrying the relevant constructs and repeated sorting of the positive population using the aforementioned method. TLR2 forms were sg-insensitive due to silent mutations at PAM region recognized by the sgRNA-Cas9 system at position R41 (AGG to CGA).

HEK293T, MC38, B16F10-FLT3L, RAW264.7, and L-929 were cultured in Dulbecco’s modified Eagle’s medium (DMEM; Gibco, 11971-025). Culture media contained 10% heat-inactivated fetal bovine serum (FBS; v/v), 2mM L-glutamine (Gibco, 25030-164) and 100 IU/ml penicillin – 100 μg/ml streptomycin (Corning, 30-001-Cl). Primary dendritic cells were cultured in R10 media: RPMI1 1640 (Gibco, 21870-076), supplemented with 10% heat-inactivated FBS, non-essential amino acids (Gibco, 11140-050), 1mM sodium pyruvate (Gibco, 11360-070), 55mM 2-mercaptoethanol (Gibco, 21985-023), 10 mM HEPES (Gibco, 15630-080), 2mM L-glutamine (Gibco, 25030-164) and 100 IU/ml penicillin and 100 μg/ml streptomycin (Corning, 30-001-Cl). Cells were maintained in a humidified incubator at 37°C and 5% CO_2._ Cells were routinely tested for Mycoplasma contamination using MycoAlert Mycoplasma Detection Kit (LONZA, LT07).

### Generation of BMDM and immortalized BMDM (iBMDM)

Bone marrow-derived macrophages (BMDM) were generated from bone marrow of 6-12 weeks old mice. Briefly, mice were euthanized with isoflurane and hindlimbs were harvested and cleaned to expose femurs and tibias. Bone marrow was flushed with complete DMEM media and cells were resuspended at 0.5x10^6^ cells/ml in DMEM (Gibco, 11971-025) media containing 20% (v/v) FBS, 30% (v/v) L-929 conditioned media, 2mM L-glutamine (Gibco, 25030-164), 100 IU/ml penicillin, and 100 μg/ml streptomycin (Corning, 30-001-Cl). Cells were differentiated for seven days in 15cm non-tissue culture-treated petri dishes with media renewal at day 4. BMDMs were harvested with cell incubation with 2mM-EDTA containing 1X DPBS at room temperature and plated on tissue culture-treated vessels one day before the experiment started. L-929 conditioned media was generated by culturing cells in T175 tissue culture-treated flasks (Corning, 431080) until complete cellular confluency was achieved. Complete media was replenished and harvested after 10 days of L929-conditioning. Media aliquots were 0.45μm-filtered (Corning, 431220), frozen down at -80°C to be thawed as needed.

Immortalized BMDM (iBMDM) were generated by transduction of BMDMs with J2 retroviruses carrying v-raf and v-myc oncogenes at days 3 and 4 as previously described ^43^. Transduced BMDM were passaged on decreased concentrations of L-929 conditioned media overtime (1^st^ week 30%, 2^nd^ week 25%, 3^rd^ week 20, 4^th^ week 15%, 5^th^ and 6^th^ week 10%, and 7^th^ and 8^th^ week 5%) until cells proliferate in DMEM complete media without L-929 conditioned media. Psi-Cre-J2 (derived from NIH3T3 cells) served as the source of J2 retrovirus.

### Tumor growth in vivo and transfer of dendritic cells

#### Isolation of CD8^+^ dendritic cells

Rubicon-deficient mice and littermate heterozygous mice were subcutaneously implanted with B16F10-FLT3L (10^7^ cells in 100 ml of 1X DPBS) to boost dendritic cells (DCs) differentiation ^42^. Enlarged spleens were isolated from tumor bearing mice 11-14 days post-implantation. Spleens were minced and digested with 1X Collagenase/Hyaluronidase (Stock 10x; StemCell Tech, 07912) at 37°C for 30 minutes. Digested spleens were filtered through 70 μm cell strainers (Fisherbrand, 22363548). After resuspension in R10 media, cells were re-filtered through 50 μm filters (Sysmex, 04-004-2327) to generate a single cell suspension. CD8^+^ DCs were isolated using CD8^+^ dendritic cell isolation kit (Miltenyi Biotec, 130-091-169), following the manufacturer protocol.

Isolated CD8^+^ DCs were cultured in R10 media supplemented with recombinant mouse FLT3L (250ng/ml; R&D Systems, 427-FL-025). CD8^+^ DCs isolated from Rubicon-proficient mice were incubated in the presence or absence of 5μM Fendiline-HCl (Tocris, 6407), CD8^+^ DCs isolated from Rubicon-deficient mice served as controls to benchmark for the effect of LAP-deficient DCs over tumor growth. After 20h, all DCs were fed killed MC38 cells for 3h and DCs were re-isolated using CD11c purification Kit (MACS, 130-125-835). MC38 were killed upon treatment with BH3-mimetic: 1μM ABT-737 (MedChem Express, HY-50907) and 1μM Mcl-1 inhibitor: S63845 (MedChem Express, HY-100741) for 2h under culture conditions. Isolated DC cells were counted, and 10 cells were intradermally injected at day 7 and day 14 after MC38 cell implantation.

#### Tumor growth

C57BL/6J female mice were subcutaneously implanted with MC38 cells (10^5^ cells in 100 μl of 1X DPBS; Gibco, 14190-144). At day 7 (palpable tumors) and day 14, tumor-bearing mice were intradermally injected with 10^6^ CD8^+^ DCs that were previously treated or not with Fendiline and fed MC38 cells (see above). Tumor-bearing mice were randomly assigned to the groups (5-6 animals per group), so each cage contained mice for different experimental groups. Operator was unaware of DC source (Rubicon-het untreated vs. Rubicon-het Fendiline-treated vs. Rubicon-KO) at the time of intradermal injection or tumor measurements. Tumor dimensions (width and length) were measured every 3-4 days (twice a week) starting at day 9. Tumor volumes were calculated using the formula 0.5 x (width^2^ x length) in two independent experiments (total of n=10-12 mice per group). Experimental endpoints were mice distress or tumor ulceration at any point or tumors volume above 2,000 mm^3^.

### Plasmids

All plasmids were validated by Sanger-sequencing before usage. Plasmid backbones, source of cDNA and cloning primers are stated in Supporting Information.

#### Lipid and LAP reporters

pMXs-Venus-Lact-C2, encodes the discoidin-type lectin domain (C2) of mouse Lactadherin/Mfge8 (residues 307-463, Uniprot: P21956) and pMXs-Venus-FVIII-C2 encodes the C2 domain of mouse Coagulation factor VIII (residues 2161-2313, Uniprot: Q06194) downstream and in-frame of mVenus (cloned MluI to NotI). These domains are reported to specifically bind phosphatidylserine ^12,32^. A cDNA library generated from mouse placenta total mRNA (Takara, 636672) served as template to amplify the domains by PCR. pMXs-Venus-LC3, was generated by subcloning full-length rat LC3B (Uniprot: Q62625) from ptfLC3 (kind gift from Dr. Tamotsu Yoshimori, Addgene#21074) downstream and in-frame of mVenus using BspHI and NotI. The pMXs-p40-phox-PX-Venus encodes the PX domain of mouse p40-phox (residues 3-148, NCF4, Uniprot: P97369) upstream and in-frame of mVenus (cloned BglII-BamHI to MluI) into the pMXs backbone. For recombinant expression, p40-phox-PX was subcloned into pGEX-4T1 and expressed as an N-terminal GST-tagged protein.

#### Rubicon constructs

Mouse Rubicon cDNA template was a generous gift from Dr. Tamotsu Yoshimori (Addgene#21636). Rubicon constructs were N-terminal tagged with either FLAG or mCherry. Rubicon 1′CCD (lacking the Beclin1-interacting region at amino acids 490-542, Uniprot: Q80U62) was generated by amplifying the N-terminal and the C-terminal fragments by PCR and later joining these using an MfeI site introduced in the amplicons. RubiconΔ645 and Rubicon1′1-644 deletions were generated by PCR. Rubicon-K718E-R719E-R721E (Rubicon-MUT, where the basic residues in the core FYVE motif were changed to acidic residues) was generated by site-directed mutagenesis. pMXs-Flag-mCherry served as a negative control for experiments involving FLAG-tagged or mCherry-tagged constructs. For recombinant expression in insect cells, full length mouse Rubicon cDNA was subcloned into pFAST-BAC-HT downstream and in-frame of a His6X N-terminal tag (BamHI-MluI) adding a C-terminal FLAG tag (MluI to NotI). Final plasmid is pFAST-BAC-HT-Rubicon-FLAG. For recombinant expression in *E. coli*, Rubicon-Δ1-644 was subcloned into pGEX-4T1 (MluI to NotI).

#### TLR2 constructs

TLR2 cDNA was a kind gift from Dr. Ruslan Medzhitov (Addgene#13083). TLR2 constructs to complement RAW264.7-TLR2-KO cells were insensitive (TLR2-sgi) to CRISPR-Cas9 activity due to silent mutation generated by site-directed-mutagenesis. This change at position R41 (AGG to CGA) eliminates the consensus for the protospacer motif (PAM) recognized by the sgRNA-Cas9 system. TLR2-sgi-K628E-R629D-K630E-K632E-K633E (called TLR2-sgi-ACID), TLR2-sgi-K628A-R629A-K630A-K632A-K633A (called TLR2-sgi-ALA), TLR2-sgi-P631H and TLR2-sgi-P681H were generated by site-directed mutagenesis of TLR2-sgi. TLR2-sgi-1′TIR and TLR2-sgi-ΔTIR-ACID were generated by PCR and lack the TIR domain (residues: 640-784, Uniprot: Q9QUN7). For recombinant expression, the complete intracellular domain of TLR2 (ID, 609-784, Uniprot: Q9QUN7), or its different mutant versions, were subcloned in different backbones. 1) For lipid arrays and PS-Beads pulldown into pGEX-4T1 (BamHI to NotI) to obtain GST-TLR2-ID, GST-TLR2-ID-ALA or GST-TLR2-ID-ACID. 2) For lipid bilayer experiments into pNIC-Bsa4 (kind gift from Dr. Tudor Moldoveanu, NdeI to NotI) to generate pNIC-His6X-TLR2-ID-FLAG and pNIC-His6X-TLR2-ID-ACID-FLAG.

#### Knockdown of component in phosphatidylserine metabolism and trafficking

First, expression of P4-ATPase flippases and co-chaperones were assessed by end-point PCR in RAW264.7 and BMDM. Primers were: *Atp11a* (cagatactgtgcaggggaaga, gacttgtgggtgtgtcgatga), *Atp11b* (gaactgccttgcagcatcg, gccattctcagtgcttcaatagt), *Atp11c* (accctcaaccgtttgtgtg, ccagaaatgggatgattgccaac), *Atp8a1* (ttagacaaggcttaccggcaa, ctttcacactcgattctgcca), *Atp8a2* (cagtgggagacatcgtgaagg, agccctgtcgtattttaaggttc), *Atp8b3* (tcggggagaaccttgaggata, tcgatggaactgctcgtacag), *Cdc50a* (caaacagcaacggctaccc, gttgttggaggtgacgaagat), *Cdc50b* (actcctccaacggcatcaag, gctcgtagtagaggtacacgg), *Gapdh* (aggtcggtgtgaacggatttg, tgtagaccatgtagttgaggtca).

Bacterial stocks of pLKO vectors expressing validated short hairpin RNA (shRNA) targeting enzymes in PS metabolism/transport were purchased from the Mission-SIGMA collection: mouse OSBPL5/ORP5 (TRCN0000105111), mouse OSBPL8/RP8 (TRCN0000105248), mouse ATP11A (TRCN0000101533), mouse ATP11C (TRCN0000101851), and mouse TMEM30A/CDC50A (TRCN0000317704). Lentiviral production and target cell transduction were performed as described below. Upon puromycin-selection, silencing was validated by western-blot or quantitative PCR (qPCR), using the primers above, if suitable antibodies were not commercially available. Silenced cells were transduced with Venus-LC3 and Venus-Lact-C2 reporters as needed.

### Transfection and transduction

HEK293T were used to produce Vesicular Stomatitis Virus-G (VSV-G) pseudotyped retrovirus and lentivirus to transduce RAW267.4 cells. Briefly, for retrovirus, HEK293T cells were co-transfected using PEI-MAX (1μg/ml; Polyscience, 324765) with VSV-G (Addgene#8454), pCL-AMPHO (Imgenex, 10046P; now in Novus) and a retroviral vector harboring the gene constructs of interest in the pMXs backbone. For lentivirus, cells were co-transfected with VSV-G, PAX2 (Addgene#12260) and a lentiviral-based plasmid (pLKO or pLenti-V2). Supernatants were collected at 48h and 72h after transfection, filtered through 0.45μm filters (Corning, 431220) and target cells were transduced twice via spinfection with the help of polybrene (8μg/ml, hexamethrine bromide, Sigma, TR1003). Cells were selected for antibiotic resistance (FLAG-tagged constructs, Puromycin 5μg/ml), or by consecutive cell sorting for the expression of fluorescent proteins (mCherry-tagged or mVenus-tagged constructs) or the re-expression of TLR2 at the cell surface (Clone T2.5, Biolegend 121810) until the cell population was homogeneous. Expression of the protein of interest was validated by western blot, immunofluorescence, and/or flow cytometry.

### Reagents

#### Compounds and drugs

Ionomycin (Cayman Chem, 11932), Fendiline-HCl (Tocris, 6407), selective PI4KIII inhibitor: GSK-A1 (Cayman Chem, 34502), BH3-mimetic: ABT-737 (MedChem Express, HY-50907); Mcl-1 inhibitor: S63845 (MedChem Express, HY-100741), mTOR inhibitor: Torin-1 (MedChem Express, HY-13003), V-ATPse inhibitor: Bafilomycin A1 (Cayman Chemical, 11038) were reconstituted as per manufactures’ datasheet and used as indicated in the figure legends. EBSS (GIBCO, 24010-043) served as starvation media.

#### Phagosome stimulation

Pam3csk4-Beads were prepared according to manufacture instructions. Briefly, ∼3μm carboxyl-polystyrene beads (Spherotech, CP-30-10) were washed twice with 1X DPBS and activated in glass tubes (Pyrex, Corning, 9826-16) for 30 min at room temperature using 10 mM Sodium Acetate [pH 5.0] buffer containing 5μg/ml 1-ethyl-3-(3-dimethylaminopropyl)carbodiimide hydrochloride (EDC, Thermofisher, 22980) as coupling agent. TLR2 ligand Pam3csk4 (Invivogen, tlrl-pms) was coupled to activated beads at a final concentration of 100μg/ml for 3h at room temperature with gentle rocking. BSA-Beads (for immunofluorescence, Spherotech; BP-30-5; ∼3μm) or Biotin-Beads (for lipidomics, Spherotech; TP-30-5; ∼3μm) served as control beads. Incubation with anti-BSA (Clone BSA-33, Sigma, B2901) or anti-Biotin (Jackson Immunoresearch Inc - JIR, 200-002-211) at 10μg/ml for 2h at 4°C rendered Ig-coupled beads to stimulate FcRs for immunofluorescence or lipidomics experiments, respectively. Zymosan particles were used as complex ligands to induce phagocytosis. Zymosan (Invivogen, tlrl-zyn), Zymosan-TexasRed (ThermoFisher, Z2843), Zymosan-Alexa594 (ThermoFisher, Z23374), Zymosan-FITC (ThermoFisher, Z2841) and Zymosan-phRODO-Green (ThermoFisher, P35365) were used as indicated in the figure legends.

#### Antibodies

Anti-Phosphatidylserine (anti-PS, Clone 1H6, EMD-Millipore 05-719), anti-Oxysterol-binding protein-related protein 8 (anti-OSBP8/ORP8; Abcam, ab99069), anti-Cell cycle control protein 50A (anti-CDC50A/TMEM30A, Sigma, AV47410), anti-Microtubule-associated protein 1 light chain B (anti-MAPLC3/LC3; for western blot: Cell Signaling Technology-CST, 2775; for immunofluorescence: MBL Int, PM036), anti-Sequestosome-1 (anti-SQSTM1/p62, Sigma, P0067), anti-phosphoinositide 3-kinase regulatory subunit 4 (anti-PIK3R4/VPS15; CST, 14580), anti-Phosphatidylinositol 3-kinase catalytic subunit type 3 (anti-PIK3C3/VPS34; Clone D9A5, CST 4263), anti-Beclin1 (CST, 3738), anti-NAPDH oxidase 2 (anti-NOX2/gp91-phox; Santa Cruz Technology, sc-130543), anti-Rubicon (Clone D9F7, CST, 8465), anti-Toll-like receptor 2 (anti-TLR2; Clone T2.5 for flow cytometry, Biolegend 121810; for western blot, Clone E1J2W, CST, 13744), anti-mCherry (Clonetech, 632543), anti-Triggering receptor expressed on myeloid cells 2 (anti-TREM2; R&D MAB17291), anti-Glutathione-S-Transferase (anti-GST, Clone GST-2; Sigma SAB4200692), anti-FLAG (Clone M2; Sigma, F1804), anti-Actin (Clone C4, HRP-conjugated; Santa Cruz, sc-47778), anti-Bovine Serum Albumin (anti-BSA, Clone BSA-33, Sigma, B2901) or anti-Biotin (JIR, 200-002-211) were used as indicated in manufactures’ datasheets.

### Immunoblotting

Cells were washed and harvested in cold 1X PBS, and then lysed for 20 min on ice. Lysis buffer was 50mM Tris-HCl [pH 7.5], 150 mM NaCl, 5mM EDTA, 1% Igepal-CA630 (v/v, Sigma, I8896); supplemented with protease (cOmplete; Roche, 11836153001) and phosphatase inhibitors (PhosSTOP; Roche, 04906837001). Cell lysates were centrifuged at 16,000xg for 10 min at 4°C, supernatants were collected, and protein concentration was quantified (BCA-based; Thermo, 23225). Same protein amount per sample was diluted with Laemmli sample buffer (4X; BioRad, 1610791) supplemented with 10% 2-betamercaptoethanol (v/v; Sigma, M3148) and 1mM dithiotreol (DTT; Sigma, D0632), boiled for 10 min at 95°C, and resolved by SDS-PAGE using Criterion XT Bis-Tris precast gels (4-12%; BioRad, 3450123/4/5) and XT-MES1X as running buffer (stock 10X; BioRad, 1610789). Proteins were transferred to 0.22μm PVDF membrane (Millipore, ISEQ00010) using tank transfer. Buffer contained 25mM Tris base, 192mM glycine and 20% methanol (v/v) in MilliQ water (final pH 8.3). Blotted membranes were blocked with 5% (w/v) milk in TBS-0.05% Tween-20 (v/v; Fisher Scientific, BP337100; TBS-t) for 1h and washed with TBS-T. Membranes were incubated overnight with antibodies in 2% (w/v) BSA, 0.01% sodium azide (w/v; Sigma, S2002) in TBS-T, washed thoroughly with TBS-T and incubated with species-specific horseradish peroxidase (HRP)-conjugated secondary antibody (Amersham, anti-mouse: NA931, anti-rabbit: NA934) in 2.5% (w/v) Milk-TBS-T. After extensive washing, membranes were developed using Clarity Western ECL substrate (Bio-Rad, 1705060). Chemiluminescence was acquired with an Odyssey-Fc device (LICOR) and ImageStudioLite (LICOR) was used as western-blotting processing software.

### Immunoprecipitation (IP)

Cell lysates were generated as described above. After protein quantification, protein amount (∼1-2mg) per sample was diluted to 0.2% Igepal-CA630 lysis buffer using buffer without detergent. Diluted lysates were subjected to immunoprecipitation with anti-Rubicon (2μl of antibody per 1mg of lysate) overnight at 4°C with rotation. Ig-complexes were precipitated with Protein A Sepharose for Fast Flow (Cytiva, 17-1279; 15μl slurry per sample) for 2h at 4°C with rotation. After extensive washes with 0.2% Igepal-CA630 lysis buffer, protein complexes were solubilized in loading buffer and analyzed by standard immunoblotting (see above). For IP of mCherry-tagged proteins, RFP-Trap agarose beads (Chromotek, rta) were used following a similar protocol. For IP of FLAG-tagged protein, anti-FLAG(M2) agarose beads (Sigma, A2220) were used. After four washes with 0.2% Igepal-CA630 lysis buffer and a final wash in TBS, FLAG-tagged proteins bound to beads were eluted using purified 3xFLAG peptide (produced in the Macromolecular Synthesis Facility at St. Jude; sequence: MDYKDHDGDYKDHDIDYKDDDDK). Eluates from three consecutive elution steps (100μg/ml 3xFLAG peptide in 1ml TBS, 20 min at 4°C with rotation) were pooled and concentrated using centrifugal filter units (Amicon, 10KDa cut-off; Millipore, Ultra-4 UFC801024 and/or Ultra-0.5 UFC501024). Same volume per sample was solubilized in loading buffer and analyzed by standard immunoblotting (see above). Alternatively, concentrated eluates from anti-FLAG IPs of phagosomes were used for VPS34 activity quantification as indicated below.

#### Immunoprecipitation for lipidomics

RAW-Rubicon-KO cells expressing mCherry-FLAG or FLAG-Rubicon (either WT or MUT) were plated in 15cm tissue culture-treated plates (∼ 25x10^6^ cells/plate, two 15cm plates per experimental point). After treatment and stimulation, cells were washed and harvested in cold 1X PBS, pelleted, and resuspended in 8ml IP-lipidomics buffer: 50mM Tris-HCl [pH 7.5], 150 mM NaCl, 1.5mM MgCl_2_ prepared in HPLC-grade water (Fisher, W5SK-4). Lysis buffer was supplemented with protease inhibitors (cOmplete; Roche 11836153001) and phosphatase inhibitors (PhosSTOP; Roche 04906837001). Cells were sonicated to obtain a homogeneous lysate: 3 cycles 10 sec, 1 min off, on ice; 50% duty cycle; Level 5 output, Sonifier450, Branson. Cell extracts were clarified at 16,000xg for 10 min at 4°C and subjected to immunoprecipitation with anti-FLAG agarose beads (50μl slurry per condition) in IP-lipidomics buffer for 3h at 4°C with gentle rocking. After three washes with 0.9% (w/v) NaCl (prepared in HPLC-grade water), beads were flashed-frozen in liquid nitrogen and kept at -80°C for further processing. An aliquot per point was run in SDS-PAGE for conventional anti-FLAG immunoblotting to evaluate amount of FLAG-tagged protein per sample and volumes for lipid extraction were normalized accordingly to process similar amounts of FLAG-tagged proteins independently of bead volume.

### Phagosome purification

Phagocytes (∼ 25x10^6^ cells/plate, two 15cm plates per experimental point) were fed with Pam3csk4-beads or control beads so that each cell engulfed 0-3 phagosomes after 45 minutes. Plates were thoroughly washed with ice-cold 1X PBS to eliminate non-phagocyted beads. Cells were harvested in ice-cold 1X PBS and mechanically homogenized in 2ml 8% (w/v) sucrose (0.25M; Sigma, S9378) using glass douncers. After homogenization, the disrupted cell solution was mixed with 62% (w/v) sucrose (1.81M, saturated solution) to reach a final sucrose concentration of ∼40% and layered onto 62% sucrose in polycarbonate centrifuge tubes (Beckman Coulter, 34058). 35% (1.02M), 25% (0.73M), and 10% (0.29M) sucrose solutions were carefully layered onto the cell solution mixtures. Sucrose-containing buffers were prepared in 3mM imidazole [pH 7.4] (Sigma, I202) and layers were 8 ml each except for the 5ml 62% layer at tube bottom. Equilibrated tubes were ultra-centrifuged at 100,000xg for 1h at 4°C using a swinging-bucket rotor (SW 32Ti, Beckman Coulter, 369650; Optima XE, Beckman Coulter). Phagosome-containing beads float at the interface between the 10% and 25% layers. Retrieved bead-containing phagosomes were washed with cold 1X PBS (or 0.9% NaCl solution for lipidomics), ultra-centrifuged at 100,000xg for 30 min at 4°C, and recovered for further processing. For binding assays, phagosomes were harvested in 1X PBS and counted using a cell counter (Cellometer, Nexcelom). For biochemical analysis, phagosomes were lysed in 1% Igepal-CA630 lysis buffer, protein concentration was determined, and equal amounts per sample were analyzed by SDS-PAGE and immunoblotting. Alternatively, phagosome lysates were subjected to anti-FLAG immunoprecipitation to analyze FLAG-Rubicon-containing complexes at the phagosome membranes or to assess VPS34-lipid kinase activity in vitro. For lipidomic assays, bead-containing phagosomes were resuspended in 0.9% NaCl (prepared in HPLC-grade water), counted to equilibrate numbers, and pelleted at 100,000xg for 20 min at 4 °C using fixed angle rotor (TLA-100 rotor, Beckman Coulter, 349481) in a benchtop ultracentrifuge (OptimaTL, Beckman Coulter). Pelleted phagosomes were transferred to lipidomics-compatible tubes in a minimal residual volume and flash-frozen in liquid nitrogen for further processing.

### In vitro analysis of VPS34 activity

Phagosomes were isolated as described above from RAW264.7-Rubicon-KO cells reconstituted with FLAG-Rubicon, FLAG-Rubicon1′CCD, or empty vector. Phagosomes were lysed in 1% Igepal-CA630 lysis buffer and FLAG-tagged Rubicon was immunoprecipitated using anti-FLAG agarose beads overnight at 4°C with rotation. FLAG-Rubicon-containing complexes were eluted with 3XFLAG peptide. Eluates and flow-through washing buffer were concentrated using centrifugal filter units and ability of fractions to generate PI3P was analyzed by competitive ELISA with Class III PI3 Kinase Kit (Echelon, K-3000), following the manufactures instructions. Aliquots were analyzed by standard immunoblotting to determine the presence of complex components.

### Analysis of Venus-LC3, LC3, p40-phox-PX-Venus, and PI3P levels by flow cytometry

Cells were plated on 12-well plates (5x10^5^ cells/well). The following day, cells were stimulated for 1h with either Zymosan-TexasRed (ZymTxR) or Zymosan-645 (Zym645). Cells were washed with 1X DPBS and harvested after 20 min of incubation with 2mM-EDTA containing 1X DPBS at room temperature. Cells were centrifugated at 300xg for 3min at 4°C and lysed with 200μg/ml Digitonin (Sigma; D141) solution in 1X DPBS for 15 min at room temperature to release free Venus-LC3. After 15 min, lysis was quenched using FACS-Buffer (1% BSA w/v, 1mM EDTA in 1X PBS), cells were centrifuged, and resuspended in FACS-Buffer. Cell fluorescence was acquired with a SP6800 Sony Spectral Analyzer and compensated values were analyzed using FlowJo.v10. When cells from different origins were analyzed (i.e., RAW264.7-VenusLC3 vs. RAW264.7-Rubicon-KO-VenusLC3 cells), values of immobilized Venus-LC3 were normalized to Venus-LC3 expression from intact cells to control for difference in reporter expression. Experiments using anti-PS blocking antibody were performed in 96-well plates and cell numbers were scaled down accordingly.

For endogenous LC3 detection by flow cytometry, Zymosan-FITC was used to stimulate the cells. Digitonin-permeabilized cell pellets were Fc-blocked (BioXCell, BE0307; final concentration: 10μg/ml) and then stained with anti-LC3 antibody (1:500; MBL Int, PM036) and anti-Rabbit-DyLight405 (1:500; JIR, 711-475-152). Incubations were performed in FACS-Buffer for 20min on ice and cells were washed with 1X PBS after primary and secondary antibody incubations.

Minor adjustments were used to detect genetically encoded p40-phox-PX-Venus by flow cytometry: LAP stimulation occurred for 30 min and digitonin solution was diluted to 50μg/ml. For endogenous PI3P detection by flow cytometry using recombinantly produced PI3P probe, 50μg/ml digitonin-treated cell pellets were Fc-blocked and stained with recombinant GST-p40-phox-PX (1μg/ml, home-made see below), followed by anti-GST (1:500, Sigma, SAB4200692) and secondary staining anti-Mouse-DyLight405 (1:500; JIR, 711-475-151). Incubations were performed in FACS-Buffer for 20min on ice and cells were washed with 1X PBS after probe, primary, and secondary antibody incubations.

### Cell imaging

#### Confocal immunofluorescence

Cells were seeded in tissue culture-treated 8-well chambered slides (mslides, IBIDI, 80826) at 50,000 cells per well. After experimental procedures, cells were fixed in 4% paraformaldehyde (v/v; PFA, Stock 16%, Electron Microscopy Sciences, 15710) for 15 min at room temperature and washed with 1X PBS. Experiments using anti-PS blocking antibody were performed in µ-Slide Angiogeneis chambers (IBIDI, 81506) and cell numbers were scaled down accordingly.

Samples that required staining were permeabilized and quenched with 0.5% (v/v) Igepal-CA630, 1% (w/v) glycine in 1X PBS for 20 min at room temperature. Fixed cells were blocked in 3% fatty acid free BSA (w/v; Sigma, 7030) in 1X PBS for 30 min at room temperature. After a 1X PBS wash, cells were incubated for 1h at room temperature with primary antibodies diluted in 2% fatty acid free BSA: anti-FLAG (1:1,000; Clone M2, Sigma, F1804), anti-Rubicon (1:500; Clone D9F7, CST, 8465), anti-LC3 (1:1,000; MBL Int, PM036), anti-VPS34 (1:500; Clone D9A5, CST 4263). Cells were washed with 1X PBS and incubated for 30min at room temperature with secondary antibodies diluted in 2% fatty acid free BSA: anti-mouse-AF647 (1:500; JIR, 715-605-151), anti-rabbit-AlexaFluor488 (1:500; JIR, 711-545-152). Stained cells were washed with 1X PBS and post-fixed in 1% (v/v) PFA in 1X PBS for 10 min at room temperature.

Images were acquired using a Marianas confocal (Intelligent Imaging Innovations, 3i) comprised of a CSU-X spinning disk, Prime95B sCMOS camera, and differential interference contrast (DIC) as well as 405, 488, 561 and 640nm laser lines were used. Alternatively, a CSU-W (Yokogawa) spinning disk to facilitate super resolution via optical reassignment (SoRA) imaging in combination with 60X 1.45NA oil objective and Prime95B camera were used. Representative pictures are shown and scale bars in confocal immunofluorescence pictures are 10μm, unless otherwise indicated.

#### Analysis of phagosome enrichment in IF images

Individual phagosomes from various cells in confocal images were analyzed by sampling a line that sections the phagosome and part of the cytosol, then fluorescence intensities on the selected channels in this region of interest (ROI) were determined. Maximum intensity at the phagosome membrane was normalized for the cytosolic expression of the probe (or the background level for the antibody staining) per ROI to determine the fluorescence enrichment at the phagosome membrane per each phagosome. Cumulative values (n>10) in various cells from different fields were analyzed and statistically compared. Value above 1 (dotted line in the graphics) indicates enrichment in the phagosome membrane.

#### Stochastic optical reconstruction microscopy (STORM)

Cells were plated on tissue-culture treated chambered coverslips (Ibidi) and allowed to engulf zymosan particles prior to fixation with 4% PFA for 10 min. Reactive groups were subsequently quenched with 20mM glycine in 1X PBS for 30 min before permeabilization with 0.1% Triton-100 for 3 min and subsequent blocking in 1X PBS buffer containing 2% BSA and 5% normal donkey serum. Anti-GFP (Rockland ImmunoChemicals; 600-401-215) and anti-mCherry (Biorbyt; orb11618) antibodies were used at 1μg/mL overnight at 4°C and were subsequently detected with STORM-appropriate secondary antibodies (Biotium; 20836 and 20811). STORM acquisition was facilitated with an N-STORM system (Nikon Instruments) as previously described^44^.

To analyze spatial distribution of single molecule data, we applied our recently developed algorithm denoted ‘normalized spatial intensity correlation (NSInC; ^45^). Briefly, 3-dimensional co-ordinates of identified single molecules are analyzed for their bi-directional association, while tested against random distribution and following correction for any edge-effect bias. The association index per field is calculated with a value of 0 representing complete spatial randomness (CSR) while values of 1 and -1 represent complete association or exclusion, respectively.

### Recombinant protein purification

#### Protein Production and Purification from bacteria

GST, GST-p40-phox-PX, GST-TLR2-Intracellular domain (ID, 589-784), GST-TLR2-ID-K628E-R629D-K630E-K632E-K633E (GST-TLR2-ID-E/D), GST-TLR2-ID-K628A-R629A-K630A-K632A-K633A (GST-TLR2-ID-ALA), GST-FLAG-Rubicon1′1-644 were recombinantly produced in *E. coli*. pGEX-4T1-based plasmids harboring the construct of interest were transformed into BL21 Star(DE3)pLysS (ThermoFisher, C602003) and plated onto Ampicillin-containing LB plates. Picked colonies were grown as 10ml volume pre-cultures in Ampicillin-containing LB broth at 37°C with vigorous shaking for overnight. Large volume Ampicillin-containing LB previously pre-warmed at 37°C were inoculated with pre-cultures. Bacteria grew at 37°C and protein induction occurred at 20°C for overnight upon addition of 0.1mM isopropyl D-1-thigalactopyranoside (IPTG; Goldbio, I2481C) to the bacterial culture at mid-log phase (OD_600nm_=0.6). Induced bacteria were pelleted by centrifugation and frozen down at -70°C for further processing. Thawed bacterial pellets were lysed in Bacterial Protein Extraction Reagent (B-PER; Thermo Fisher Scientific, 90079) supplemented with 100μg/ml lysozyme (Thermo Fisher Scientific, 90082) and 5U/ml DNase-I (Thermo Fisher Scientific, 90083) for 30 min at room temperature with gently rocking. Bacterial lysates were clarified by centrifugation at 30,000xg for 15 min at 4°C, followed by subsequent centrifugation at 30,000xg for 30 min at 4°C. Supernatants containing soluble recombinant protein were pulled-down with Glutathione (GSH) Sepharose 4FastFlow (GE Healthcare, 17-5132) for 3h at 4°C with gently rocking. Beads were extensively washed with 0.2% Igepal-CA630 TBS (x3), followed by two washes with 100mM Tris-HCl [pH8.0],150mM NaCl, 5mM EDTA Buffer. GST-tagged recombinant protein was eluted by three sequential incubations of GSH-beads with 50mM reduced L-GSH (SIGMA, G4251) in 100mM Tris-HCl [pH8.0],150mM NaCl, 5mM EDTA buffer for 30 min at 4°C with rocking. The pooled eluates were concentrated using centrifugal filter units (Amicon, 10KDa cut-off; Millipore, Ultra-15 UFC901024 and Ultra-0.5 UFC501024). Purification yield and protein integrity were analyzed by SDS-PAGE followed by Coomassie staining according to manufacturer instructions (PageBlue Protein Staining Solution; Thermo Fisher Scientific, 24620). Protein concentration was determined with NanoDrop measurements.

His6X-TLR2-ID-FLAG and His6X-TLR2-ID-K628E-R629D-K630E-K632E-K633E-FLAG (His6X-TLR2-ID-ACID-FLAG) were recombinantly produced in *E. coli*. pNIC-based plasmids harboring the construct of interest were transformed into BL21 Star(DE3)pLysS (ThermoFisher, C602003). Bacteria were induced as previously described. Frozen induced-bacterial pellets were resuspended in lysis buffer containing 20mM Tris-HCl [pH 8.0], 300mM NaCl, 1mM PMSF, 1mM AEBSF, 1X protease inhibitor cocktail and 10% glycerol at 4°C using 10ml per gram of bacterial pellet. Lysis was performed by microfluidizer at 4°C, lysates were processed twice to insure complete lysis. Lysates were clarified 18,500xg for 1h at 4°C (Avanti JXN-26, rotor JA-25.50) and subjected to metal affinity chromatography (50ml 50% Ni-NTA slurry per condition). Equilibration buffer for the column was 20mM Tris-HCl [pH 8.0], 300mM NaCl, and 10% glycerol. After gravity flow, the column was washed with four bed volumes of equilibration buffer containing 5mM, 10mM, 20mM and 30mM imidazole, respectively. Final elution proceeded with four bed volume of equilibration buffer containing 400mM imidazole. Fractions were analyzed by SDS-PAGE, followed by gel staining and elution fractions containing the protein of interest (5mM, 10mM and 20mM) were pooled and subjected to size exclusion chromatography. HiLoad 26/600 Superdex 75pg (MW range: ∼3,000 to ∼ 70,000 KDa; Cytiva Lifescience, 28989334) was the column and equilibration buffer contained 20mM Tris-HCl [pH 8.0], 50mM NaCl. Size exclusion chromatography was performed on AKTA-PURE at 4°C and recombinant proteins eluted as a single peak: 163.49ml for WT version and 156.98ml for ACID version. Fractions containing the protein of interest were pooled and concentrated using centrifugal filter units (Amicon, 10KDa cut-off; Millipore, Ultra-4 UFC801024 and/or Ultra-0.5 UFC501024). Final protein concentration was determined by nanodrop measurements.

#### Protein production in insect cells

Full length mouse Rubicon cDNA was FLAG-tagged at the C-terminus and subcloned into pFAST-BAC-HT. Double termini tagging (His6X-tag at the N-terminus and FLAG-tag at the C-terminus) allowed tandem purification to ensure full-length protein to be recovered. His6X-Rubicon-FLAG was expressed and purified from insect cells in the Protein Production Facility at St. Jude Children’s Research Hospital. Briefly, pFAST-BAC-HT-Rubicon-FLAG was transformed into DH10Bac E.coli (Vendor?, Cat No.?) and used to generate bacmid DNA that was used to transfect Sf9 insect cells (Vendor?, Cat No.?) using serum-free media. Transfected Sf9 insect cells generated, by homologous recombinant, baculovirus harboring His6X-Rubicon-FLAG. Baculovirus-containing supernatant from transfected cells was used to infect Sf9 that amplified viral stock to serial infection to finally infect 3 liter of Sf9 cells. After 72h, infected cells were harvested and frozen at -20°C.

The frozen cell pellets were lysed using a microfluidizer in buffer containing 50 mM Tris-HCl, pH 8.0, 500mM NaCl, 10% glycerol and the lysate centrifuged at 20,000 rpm for 1h at 4°C. The supernatant was 0.2 µM filtered and was incubated overnight at 4°C with 5 mL of Ni-NTA beads (Qiagen). XXX

### Lipid binding assays

#### Protein-lipid overlays – Lipid strips

Lipid binding capacity of recombinant proteins or anti-PS antibody were analyzed using lipid arrays (Echelon Biosciences Inc; Membrane lipid Strips, P6002, or PIP Strips P6001) as directed by manufacture protocols. Briefly, membranes spotted with different lipid species were blocked with 3% fatty acid free BSA (w/v; Sigma, 7030) in 0.1% (v/v) Tween-20 1X PBS (PBS-T) for 1h at room temperature with gentle rocking. Recombinant protein was diluted to 500ng/ml in blocking buffer and incubated for 1h at room temperature with gentle rocking. Primary anti-GST (1μg/ml), and secondary anti-mouse-HRP (1:5,000) antibodies revealed protein-lipid interaction. Extensive washes with PBS-T were performed between probe, primary and secondary antibody incubations. Protein-lipid overlay arrays were developed as conventional immunoblot and ImageStudioLite (LICOR) was used to performed densitometry analysis of chemiluminescence spots. Values were normalized to the value of blank control per image. To analyze the selectivity of the anti-PS antibody (Clone 1H6, EMD-Millipore 05-719) the primary antibody was added after blocking at 1μg/ml, rest of steps remained the same.

#### PS-bead pulldown

Recombinant mTLR2-ID wild-type, ALA-mutant, or ACID-mutant purified from *E. coli* (see above) were analyzed side-by-side for their binding-capacity to PS-conjugated beads (Echelon Biosciences Inc, P-B0PS) or control beads, as directed by manufacture protocols. Briefly, 15 μl of slurry beads per point were washed twice in binding buffer: 10mM HEPES [pH 7.4], 150mM NaCl, 0.25% (v/v) Igepal-CA630; then 10 μg of recombinant protein was incubated for 3h at 4°C with rotation. After 4 washes with binding buffer, conjugated proteins were subjected to conventional immunoblotting for anti-GST.

#### Competition assays on isolated Pam3csk4-bead-containing phagosomes

Pam3csk4-beads containing phagosomes were isolated from RAW264.7-Rubicon-KO cells as described above. Same number of phagosomes were incubated in half-area 96 well-plates with the PS-binding probe bovine Lactadherin-FITC (BLAC-FITC, Haemtech Co, now Prolytix), recombinantly produced GST-FLAG-RubiconΔ645 or their combination at different concentrations for overnight at 4°C with gentle rocking. After PFA fixation, phagosomes were permeabilized and quenched with 0.5% (v/v) Igepal-CA630, 1% (w/v) glycine in 1X PBS for 20 min at room temperature, blocked with 3% (w/v) BSA in 1X PBS, stained with primary (anti-FLAG) and secondary antibodies (anti-mouseCy3; JIR, 115-165-166) and post-fix with 1% (v/v) PFA for 10min. Phagosomes were imaged by confocal microscopy and green and red signals were quantify per phagosome.

### Planar glass-supported lipid bilayers Peptides

All intracellular domains contain six-histidine tag (HisX6) at their N-termini and a PEG-Biotin moiety (or a FLAG-tag) at their C-termini, sequences can be found in Extended Table 1. His6X forced the proper orientation of the cytosolic tail in the planar glass-supported lipid bilayer, the biotin moiety (or the FLAG-tag) allowed streptavidin-based (or anti-FLAG antibody) intracellular domain manipulations. Intracellular domains were based on Uniprot annotation: mouseCD16 (P08508, residues 236-261), humanCD16 (P08637, residues 230-254), humanTIM4 (Q96H15, residues 336-378), and mouseTLR2 (Q9QUN7, residues 606-784). Solubilization after lyophilization of mouseTIM4 (Q6U7R4, residues 301-343) intracellular domain was ineffective, and therefore human TIM4 was used.

TLR2-ID was recombinantly produced in bacteria (see above). Small intracellular domains for mCD16, hCD16, hTIM4 and mTIM1 (hereafter cytosolic tail) were synthesized as peptides in the Macromolecular Synthesis Facility at St. Jude using a SymphonyX peptide synthesizer (Gyros Protein Technologies) and standard Fmoc chemistry. Peptides contained a C-term PEG-Biotin and were synthesized using a preloaded PEG-Biotin resin (Sigma-Aldrich). Peptides were cleaved from the resin using TFA / Water / Thioanisole / Triisopropylsilane / Phenol / Ethanedithiol – 82.5/0.5/0.5/0.25/0.25/0.25 and precipitated into cold diethyl ether, followed by centrifugation and lyophilization. Crude peptides were analyzed for purity using a Waters Alliance HPLC system fitted with a 2489 UV-visible detector and a 2475 fluorescence detector. Mass spec of peptides was confirmed using a Bruker Microflex LRF.

#### Lipid bilayers

All lipids were purchased from Avanti Polar Lipids and planar glass-supported lipid bilayer were prepared as previously described ^46^. Briefly, a lipid mixture containing 30% 16:0-18:1 PC (POPC, 850457), 50%PE (PE, 792518), 10% 18:1 DGS-Ni-NTA (790404), 10% PS (PS, 940037) and 1% TopFluor PS (810283) was mixed in chloroform, dried under vacuum and resuspended in 10ml of 1X PBS. After lipid extrusion using a mini-Extruder (Avanti Polar Lipids), 50μl of liposome solution was added to 1ml of bilayer buffer (20mM HEPES, 50mM NaCl) and 150μl of solution was deposited on a glass slide affixed to a flow cell (Ibidi; catalog 80608) previously cleaned with piranha solution (1:1 mixture 30% H_2_O_2_ and 96% H_2_SO_4_). Lipid-bilayer was incubated for 5min at room temperature for equilibration, excess of lipid was washed off with 2 washes of 1ml bilayer buffer and sample lanes were imaged for TopFluor-PS clustering to determine baseline clustering of PS in the lipid bilayer (Lipids Only in the panels). N-terminal His6x-tag peptides were diluted X100 in 1X PBS and a planar glass-supported lipid bilayer was incubated with 500ml of peptide solution ∼5-10μM for 15min at room temperature. After wash, images were acquired to assess the effect of peptides on TopFluor-PS clustering. Manipulations of peptide/protein on planar glass-supported lipid bilayer was achieved by incubation in high salt concentration (20mM HEPES 150mM NaCl, for 10 min at room temperature) or non-labelable streptavidin (10μg/mL, Sigma-Aldrich, Cat No. 189730). For His6x-TLR2-ID-FLAG, anti-FLAG antibody (10μg/mL, Biolegend, Cat No. 637301) was used instead. Images were acquired with a Marianas spinning disk microscope (Intelligent Imaging Innovations) equipped with SoRa CSU-W (Yokogawa), Prime 95B sCMOS camera (Photometrics) and 1.45 NA 100X oil objective. Images were acquired and analyzed using Slidebook software version 6.0.24 (Intelligent Imaging Innovations).

### Receptor recycling assay

TREM2 recycling by LANDO was analyzed as previously described ^2^. Briefly, cells were plated on 4-well chambered slides (μslides, IBIDI, 80426). The next day cells were blocked with 10% normal donkey serum in DMEM (v/v; Sigma, S30-M) for 15 min at 37°C, followed by incubation with anti-TREM2 (1:100; R&D, MAB17291) in 1% donkey serum and 5% mouse serum in DMEM for 1h at 37°C. Antibody-containing medium was aspirated, cells were acid stripped with cold DMEM (pH 2.0) and washed twice with cold 1X DPBS to remove cell-surface antibody. Cells were then re-incubated in 10% donkey serum in DMEM for 1 hour at 37°C to allow recycling of the internalized receptor-antibody complexes at the cell surface. These were labelled with secondary Alexa Fluor 594-antibody (1:500; Thermo Fisher Scientific, A-21209) in 1% donkey-serum in DMEM for 1 hour at 37°C. Cells were then acid stripped, washed with cold 1X DPBS and fixed in 4% (v/v) PFA for 15 min at room temperature. Cell-permeable Hoechst dye was added to label nuclei. Images were acquired on a Marianis spinning disk confocal microscope (Intelligent Imaging Innovations, 3i) equipped with an EMCCD camera. Image analysis including all quantification was performed using the software Slidebook 6 (3i). Quantification of recycled TREM2 receptors was performed by calculating the sum of the intracellular fluorescent signal divided by the total number of cells.

### Yeast killing assay

*Saccharomyces cerevisiae* was purchased from ATCC (Cat No. 201389) and cultured at 30°C in Yeast Peptone Dextrose (YPD) agar plates or liquid broth (Sigma-Aldrich, Y1500, Y1375). Macrophage capacity for *S. cerevisiae* killing was analyzed as previously described ^5^. Briefly, macrophages were plated in 12-well plates at 5x10^5^ cells/well, triplicates per time point were used. Yeast cells from overnight YPD-liquid culture were washed three times with 1X DPBS, number of yeast were estimated based on OD600nm turbidity, and then added to macrophages at 1:1 ratio. After 1h, wells were extensively washed with 1X DPBS. One set of triplicates served as baseline value (time 0h) and macrophages were lysed by osmotic shock upon incubation with MilliQ-H_2_O for 10 min at room temperature, then serially diluted to 1:10,000 and plated onto YPD-agar plates in duplicate per well. The remaining wells were maintained in complete medium and processed as described above at the indicated time points. YPD-agar plates were incubated at 30°C for 24h or until colonies were clearly visible and yeast colonies were counted. Yeast killing capacity was normalized to baseline values per cell line to account for differences in plating or phagocytosis for each cell line. Final values were represented as percentage of viable yeast.

Effects of GSK-A1 and Fendiline-HCl on yeast growth in YPD liquid culture was determined by assessing yeast culture turbidity at OD60nm with spectrophotometer overtime.

### Phagosome acidification assays

Cells were plated onto 48well plates (75,000 cells per well) using 6-8 replicates per condition. Following day, after treatments, cells were fed the pH sensitive probe zymosan-pHrodo-Green particles (0.2 μl per well prepared as a mastermix) to assess acidification of phagosomes overtime. The probe remains colorless at neutral pH and turns green upon acidification. Images were acquired every 60min using Incucyte (Essen Biosciences) and manufacture’s image software allowed cell segmentation and fluorescent quantification overtime. Experiments using different cell lines showed comparable values of confluency overtime and similar levels of phagocytosis were confirmed by flow cytometer in parallel assays.

### Lipidomics analyses

#### Lipids extraction

Phagosomes, cells, or IP beads were used for lipidomic analysis; samples were processed as described above. Equal numbers (3×10^6^) of macrophages, (∼5x10^6^) phagosomes or IP beads were washed with ice-cold 1X DPBS, flash-frozen in liquid nitrogen and then stored at –80°C until samples were processed for extraction of total lipids. A modified Folch extraction procedure ^47^ was used for the extraction of total lipids from purified sample. Briefly, 1 ml of chloroform-methanol (2:1, v/v) was added to the cells or beads and mixed by vortexing. Next, 200 µl of saline was added, and the tubes were mixed for 30 sec in a Bead Ruptor Elite (OMNI International) for 30 sec at 8 m/s. The homogenate was incubated at room temperature for 30 sec and then centrifuged for 10 min at 21,000xg at 4°C. After centrifugation, the lower organic-phase layer was transferred to a new tube and evaporated to dryness under a stream of liquid nitrogen. The dried lipid extracts were thoroughly dissolved with 30 µl of chloroform-methanol (2:1, v/v), transferred to autosampler vials and analyzed by LC-MS/MS (10 µl per injection).

#### LC-MS lipid profiling

LC separations were performed with a Vanquish Horizon UHPLC (Thermo Fisher Scientific) using stepped-gradient conditions as follows: 0–4.5 min, 45 to 60%, B; 4.5–5 min, 60 to 70%, B; 5–8 min, 70%, B; 8–19 min, 70 to 75%, B; 19–20 min, 75 to 90%, B; 20–33 min, 90 to 95%, B; 33–34 min, 90 to 100%, B; 34–39 min, 100%, B; 39–40 min, 100 to 45%, B; 40–45 min, 45%, B. Mobile phase A was water/acetonitrile (60:40, v/v) and mobile phase B was IPA/acetonitrile (90:10, v/v); both A and B contained 10 mM ammonium acetate. The column used was a Thermo Fisher Scientific Accucore C30 (2.1 mm × 250 mm, 2.6 μm) operated at 50°C. The flow rate was 250 μl/min and the injection volume was 10 μl. A Thermo Fisher Scientific Q Exactive hybrid quadrupole-Orbitrap mass spectrometer (QE-MS) equipped with a HESI-II probe was employed as detector. For each sample, two chromatographic runs were carried out subsequently, and separate data were acquired for negative and positive ions. The QE-MS was operated using a data-dependent LC-MS/MS method (Top-15 dd-MS^2^) for both positive and negative ion modes. The mass spectrometer was operated at a resolution of 140,000 (FWHM, at m/z 200), AGC targeted of 1 × 10^6^, and max injection time 80 msec. The instrument’s operating conditions were: scan range 100–1,500 m/z; sheath gas flow 45; aux gas flow 8; sweep gas 2; spray voltage 3.6 kV for positive mode and 2.5 kV for negative mode; capillary temperature equal to 320 °C; S-lenses RF level 50; aux gas heater equal to 320°C. For the Top-15 dd-MS^2^ conditions a resolution of 35,000 was used, AGC targeted of 1 × 10^5^, max injection time 50 msec, MS^2^ isolation width 1.0 m/z, NCE 35.

#### Data processing

The Thermo Fisher Scientific LipidSearch software (version 4.2) was used for identification and relative quantification of lipids with the following parameters: precursor and product ion mass tolerance of ± 5 ppm; main adducts search (M+H, M-H, M+NH_4_, M+CH3COO, M+2H, M-2H, M+Na, M+K) for all precursor ions. All lipid sub-classes were searched within for the major lipid classes (phospholipids, sphingolipids, glycerolipids and neutral lipids). All individual data files were searched for product ion MS/MS spectra of all lipid precursor ions. The MS/MS predicted fragmented ions for all precursor adducts were measured within 5 ppm of mass tolerance. The product ions that matched the predicted fragment ions within 5 ppm of mass tolerance were used to calculate a match-score, and those candidates providing the highest quality match were determined and used for the identification of lipid molecules, and the peak areas integrated to generate chromatographic data for semi-quantitative analyses. Next, the resulting data from search results was used to perform alignments across the experimental groups under the following Alignment setup: ExpType LC-MS; Alignment Method Mean; R.T. Tolerance 0.25 min; Calculate unassigned peak area On; Filter Type New filter; Toprank filter On; Main Node Filter All isomer peaks; m-Score Threshold 5.0; c-Score Threshold 2.0; ID Quality filter A, B, C and D.

The sum of all peak areas was taken as total lipid content per sample and individuals lipid values were normalized to total lipid content to account for slight differences in lipid extraction or data acquitsion. The data obtained from LipidSearch alignments was exported to Excel, formatted to comma-separated value (CSV) files, normalized as indicated above and then imported into MetaboAnalyst 5.0 for multivariate data analysis. The peak areas were normalized using the parameters for sample normalization sum, data transformation log_10_ and data scaling range. After the normalization in MetaboAnalyst, statistical analysis using multiparametric ANOVA, Partial Least-Squares Discriminant Analysis (PLS-DA) and Heatmaps using the default clustering algorithms were made to interrogate the lipidomics data looking for significances of individual molecules and lipid classes.

#### Software

FlowJo v10 was used for analysis of flow cytometry and GraphPad was used for data statistical analysis and data visualization. Other software packages have been indicated in specific methods sections. Cartoons were generated using BioRender.

## Acknowledgments

We thank current and past members from Dr. Doug Green’s lab and department of immunology in St. Jude for comments, reagents, protocols sharing, and support. Mao Yang and Xiaofei Wang helped with mouse work and tumor measurements. Patrick Rodrigues (Macromolecular Synthesis Facility at St. Jude) synthesized the peptides. Richard Heath, Youming Shao, Vibhor Mishra, Muralidhar Reddivari, Terry Coop, and Rosario Mosca (Protein Production Facility at St. Jude) helped with recombinant protein production.

## Author contributions

Conceptualization: EBR, DRG.

Methodology: EBR, CSG, GP, LM, ZL, DRG.

Investigation: EBR, CSG, GP, LM, ZL.

Writing - Original Draft: EBR, DRG.

Writing - Review & Editing: EBR, DRG, CSG, GP, LM, ZL.

Visualization: EBR.

Supervision: DRG.

Funding acquisition: EBR, DRG.

## Data and materials availability

All data are available in the main text, extended data, or the supplementary materials. Source data for lipidomics experiments is available upon request to corresponding author and will be deposited in the relevant repository upon manuscript acceptance. Plasmids used in this study will be deposited in Addgene or are commercially available.

Correspondence and requests for materials should be addressed to Douglas R. Green

