## Extended Figures for "Phosphatidylserine clustering by membrane receptors triggers LC3-associated phagocytosis"

Extended Data Figure 1. Boada-Romero et al.

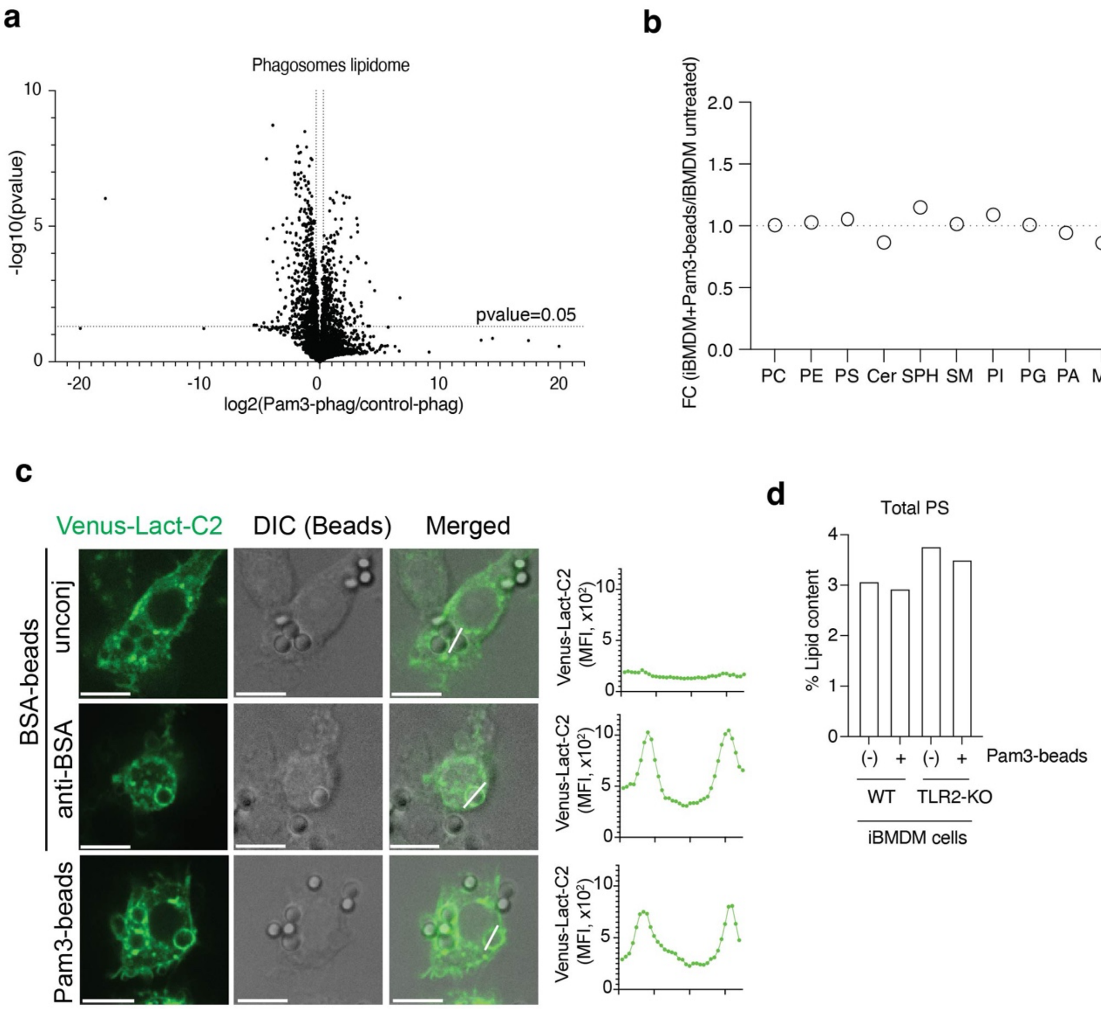


**Extended Data Figure 1: Receptor-mediated PS enrichment in phagosome membranes.** (**a**) Volcano plots showing all lipid species in Pam3csk4-beads (Pam3-phag) relative to uncoupled-beads (control-phag). Horizontal line marks statistical significance (T-test, p-value=0.05; -log10(p-value)=1.3). (**b**) Cellular lipid levels of iBMDM fed Pam3csk4-beads (Pam3-beads) relative to untreated iBMDM. Lipid species determined by lipidomics were aggregated per lipid class, shown is fold change (FC). PC: phosphatidylcholine; PE: phosphatidylethanolamine; PS: phosphatidylserine; Cer: ceramide; SPH: sphingosine; SM: sphingomyelin ; PI: phosphatidylinositol; PG: phosphatidylglycerol; PA: phosphatidyl acid; MG: monoacyl-glycerol; DG: diacyl-glycerol; TG: triacyl-glycerol; (**c**) RAW264.7 cells stably expressing the PS-probe Venus-LACT-C2 were fed BSA-beads, BSA-beads coupled with anti-BSA antibody (anti-BSA), or Pam3csk4-beads (Pam3-beads) for 30min. Representative images and profiles of MFI of Venus channels across the region-of-interest (white lines in images) are shown. (**d**) Wild-type (WT) or TLR2-KO iBMDM fed Pam3csk4-beads (Pam3-beads), as indicated, and PS content were determined by lipidomics. Shown is the mean of 5 replicates to total lipid content per each sample in one representative experiment (n=2).

Extended Data Figure 2. Boada-Romero et al.

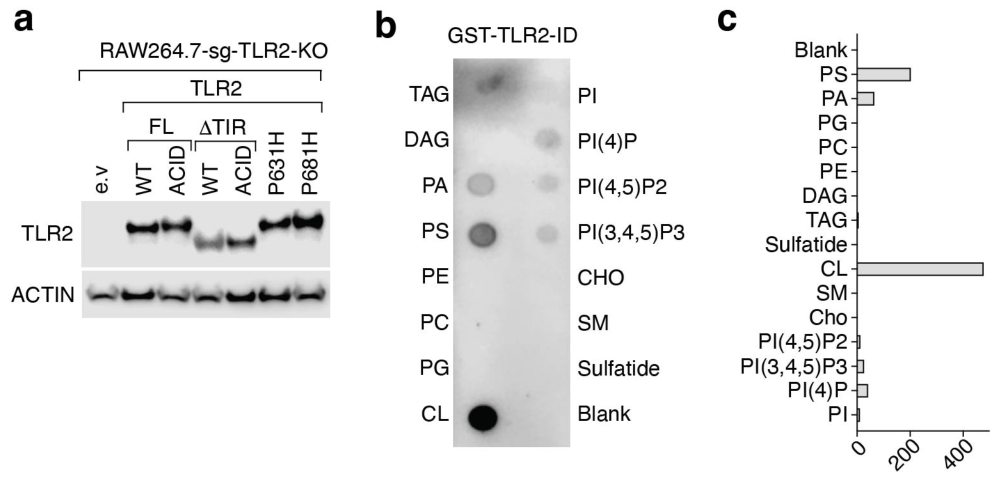


**Extended Data Figure 2: The TLR2 intracellular domain binds to phosphatidylserine**. (**a**) RAW264.7-sg-TLR2-KO cells were restored to express full length (FL) TLR2 or lacking the TIR domain (TLR2ΔTIR), in either wild-type (WT), K628E-R629D-K630E-K632E-K633E (ACID), P631H (corresponding to human SNP rs5743704), or P681H mutant, as indicated. Transduction with empty vector (e.v.) serves as a negative control. (**b**-**c**) Lipid strip showing the binding specificity of TLR2 intracellular domain (TLR2-ID) recombinantly produced in bacteria. (b) Immunoblot and (c) levels were quantified by densitometry and normalized to the value of *blank* (dotting buffer). PS: phosphatidylserine; PA: phosphatidyl acid; PG: phosphatidylglycerol; PC: phosphatidylcholine; PE: phosphatidylethanolamine; DAG: diacyl-glycerol; TAG: triacyl-glycerol; CL: cardiolipin; SM: sphingomyelin; CHO: cholesterol; PI: phosphatidylinositol.

Extended Data Figure 3. Boada-Romero et al.

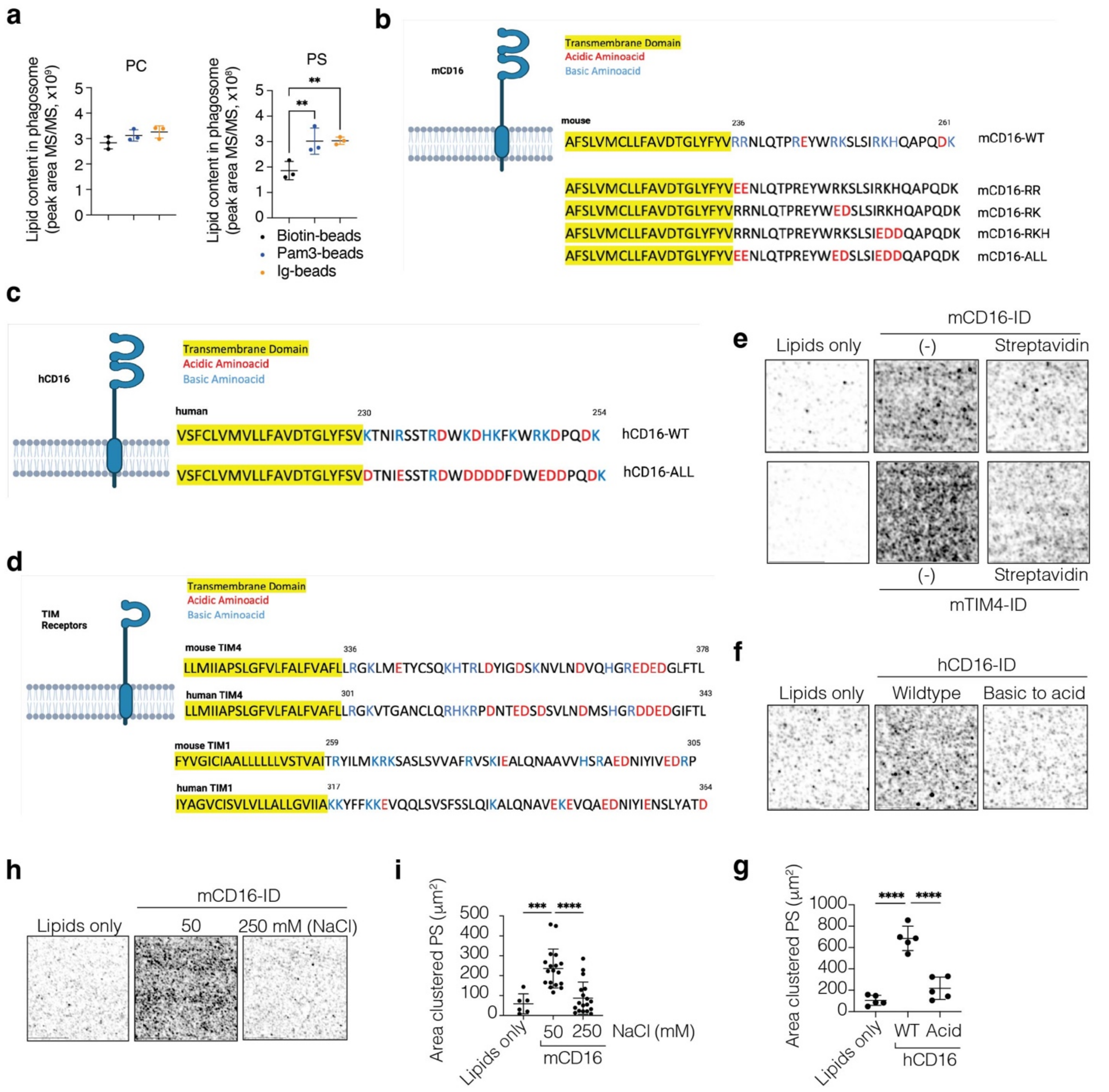


**Extended Data Figure 3: Basic patches in the intracellular domains of LAP-inducing receptors cluster PS.** (**a**) Aggregated MS/MS values per lipid species in phagosomes containing Pam3csk4-beads (Pam3-beads) or Ig-coupled beads (Ig-beads) relative to phagosomes containing control-beads (biotin-beads) isolated from immortalized bone marrow derived macrophages (iBMDM). Lipid species determined by lipidomics were aggregated per lipid class. Data are means ± SD of 3 biological replicates. Representative of n=2 independent experiments. PC, phosphatidylcholine; PS, phosphatidylserine. (**b-d**) Scheme of CD16 (mouse, Uniprot: P08508;

human, Uniprot: P08637), and TIM4 (mouse, Uniprot: Q6U7R4; human, Uniprot: Q96H15) depicted at the plasma membrane. Transmembrane domains are highlighted in yellow; acidic and basic amino acid are red and blue, respectively. Numbers indicate amino acid position. (b) Basic to acidic mutants are indicated for mCD16 and (c) hCD16. mCD16-RR: R236E-R237E; mCD16-RK: R247E-K248D; mCD16-RKK: R253E-K254D-K255D. hCD16-ACID indicate all previous basic residues mutated to acidic. (**e-i**) Glass-supported lipid bilayers resembling plasma membrane composition were incubated with cytosolic tails of mCD16 (e, h), hCD16 (g) or hTIM4 (e) as indicated, and top-Fluor-PS clustering was determined by immunofluorescence. (e, f, h) Representative images or (g, i) area of clustered PS quantified in different fields from n=2 independent experiments. ***$P$ < 0.001, ****$P$ < 0.0001 by two-sided Student’s t test.

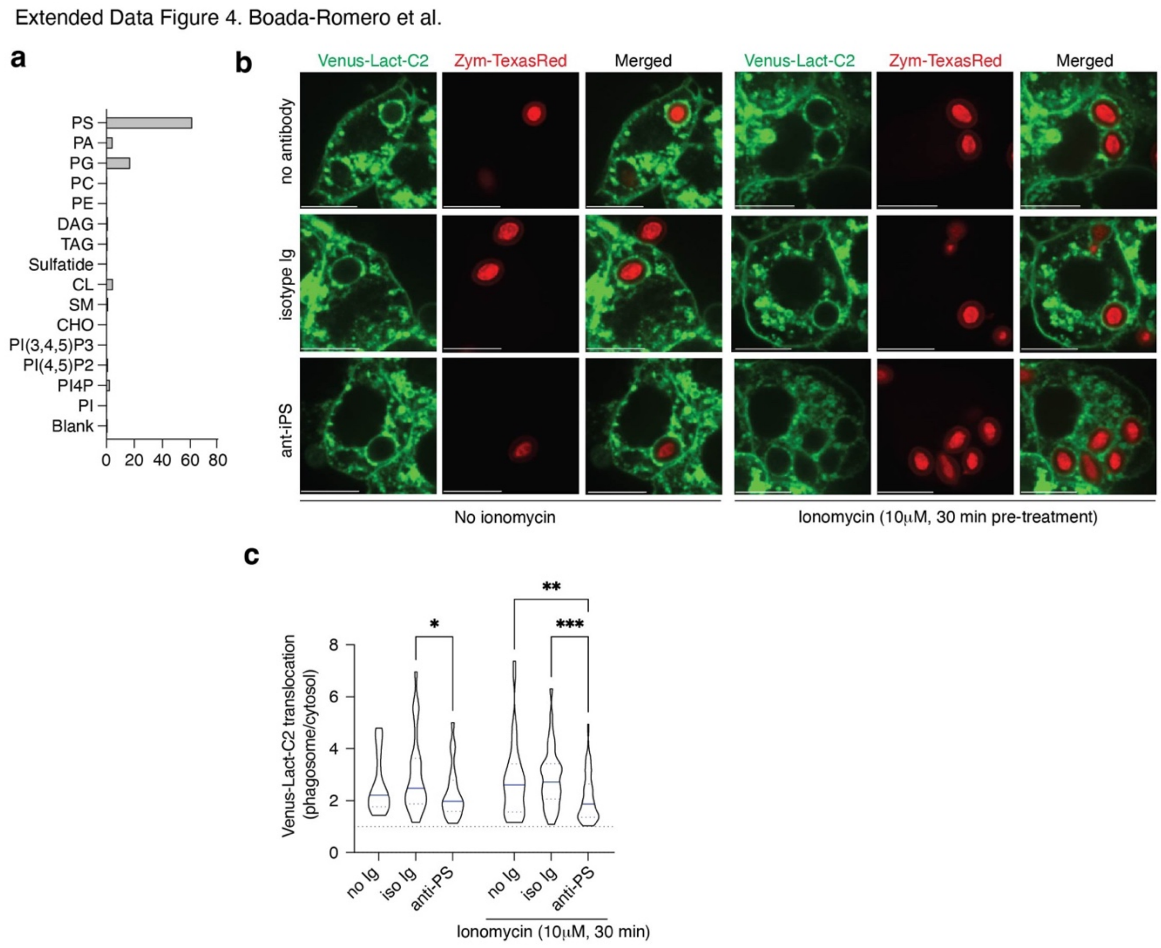


**Extended Data Figure 4: PS scrambling combined with anti-PS antibody treatment blocks PS enrichment**. (**a**) Lipid strip showing the binding specificity of anti-PS antibody. Levels were quantified by densitometry and normalized to the value of *blank* (dotting buffer). PS: phosphatidylserine; PA: phosphatidyl acid; PG: phosphatidylglycerol; PC: phosphatidylcholine; PE: phosphatidylethanolamine; DAG: diacyl-glycerol; TAG: triacyl-glycerol; CL: cardiolipin; SM: sphingomyelin; CHO: cholesterol; PI: phosphatidylinositol. (**b-c**) RAW264.7 cells stably expressing the PS-probe Venus-LACT-C2 were pre-treated as indicated (30 min) and fed Zymosan-TexasRed (Zym-TexasRed, 30 min). (b) Representative confocal images and (c) violin-plots showing the enrichment of the PS-probe at the phagosome membranes relative to the cytosol (n>18 phagosomes). Representative of n=2 independent experiments. Violin plots with median and upper and lower quartiles. **P* < 0.05, ***P* < 0.01, ****P* < 0.001 by ANOVA (pairwise comparations Fisher's LSD).

Extended Data Figure 5. Boada-Romero et al.

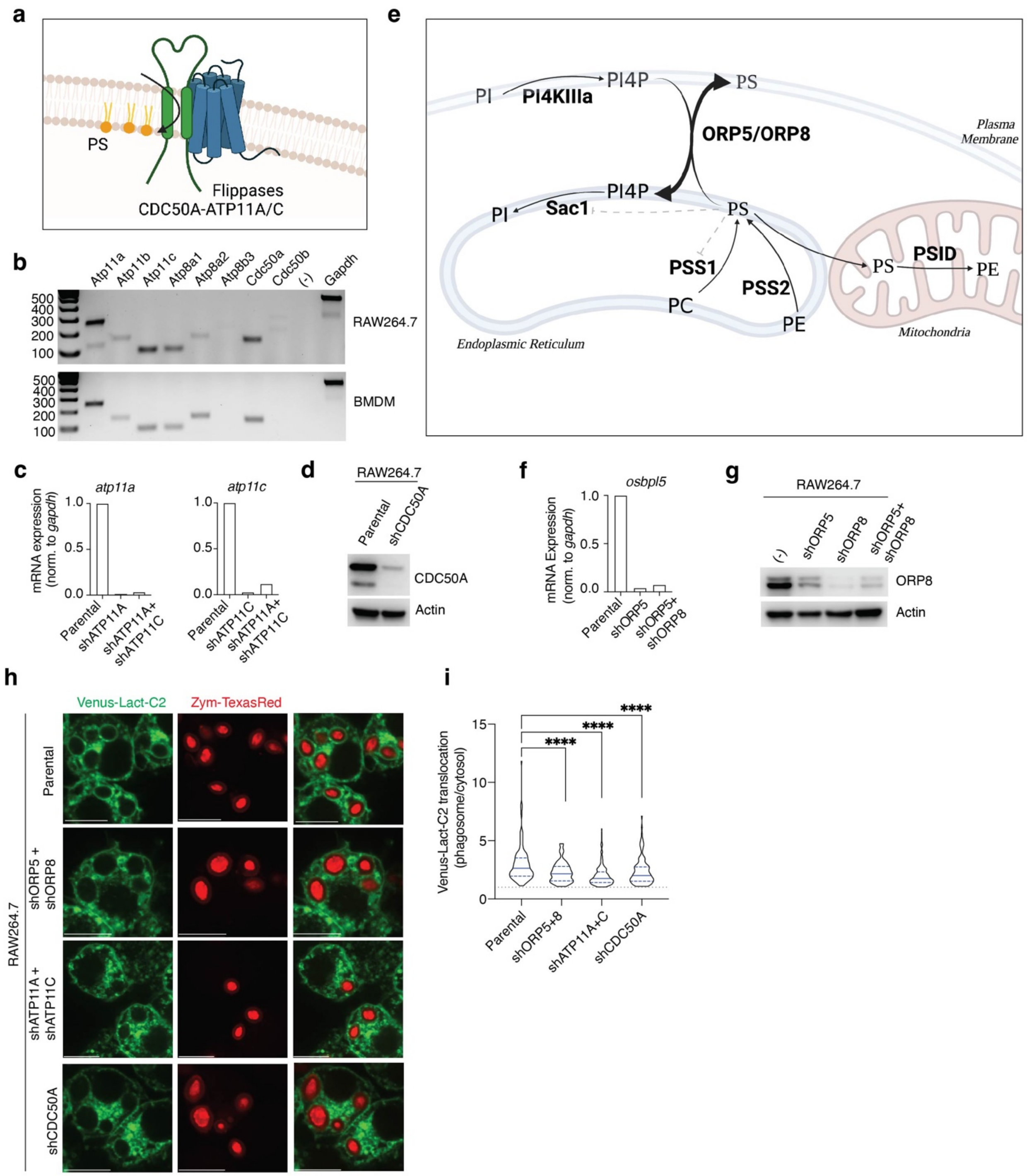


**Extended Data Figure 5: Additional strategies to limit PS enrichment in phagosomes. (a)** Scheme showing protein complexes for phosphatidylserine (PS) translocation to ensure the maintenance of PS asymmetric distribution at the plasma membrane. P4-ATPase flippase complexes are composed by phospholipid-transporter ATPase 11a (ATP11A) or ATP11C together with the co-chaperone CDC50A/TMEM30A. (**b**) Gene expression of PS P4-ATPase flippases and co-chaperones in RAW264.7 and bone marrow derived macrophage (BMDM). Higher expression is detected for Atp11a and Atp11c and co-chaperone Cdc50a. (**c, d**) Silencing efficiency in RAW264.7 cells transduced with lentivirus harboring a short-hairpin (sh) as indicated. (c) Quantitative PCR determination of *atp11*a and *atp11c* relative to *gapdh* and normalized to the levels in RAW264.7 parental cells. (d) Immunoblotting for CDC50A protein levels in parental and shCDC50A cells. (**e**) Scheme showing PS metabolism and transport in the cell. PS is synthetized in the endoplasmic reticulum (ER) by headgroup exchange via phosphatidylserine synthase-1 (PSS1) using phosphatidylcholine (PC) and via PSS2 using phosphatidylethanolamine (PE). PS is catabolized by mitochondrial phosphatidylserine decarboxylase (PSID) to produce PE. PS synthetized in the ER can be transported to the plasma membrane by Oxysterol-binding protein-related protein 5 (ORP5)/ORP8 heterodimer as an antiport using phosphatidylinositol 4-phosphate (PI4P). PI4P is in turn generated by phosphorylation of PI via phosphatidylinositol 4-kinase alpha (PI4KIIIa) and PI4P can be dephosphorylated to PI by phosphatidylinositol-3-phosphatase SAC1 in the ER. (**f, g**) Silencing efficiency in RAW264.7 cells transduced with lentivirus harboring shRNA as indicated. (**f**) Quantitative PCR determination of *osbpl5* mRNA levels relative to *gapdh* and normalized to the levels in parental cells. (**g**) Immunoblotting for ORP8 protein levels. (**h, i**) RAW264.7 stably expressing the PS-probe Venus-LACT-C2 and harboring the indicated shRNA were fed Zymosan-TexasRed (Zym-TexasRed, 30 min). (h) Representative confocal images and (i) violin-plots showing the enrichment of PS-probe at the phagosome membranes relative to the cytosol (n>70 phagosomes) in two independent experiments. ****$P < 0.0001$ by ANOVA test.

Extended Data Figure 6. Boada-Romero et al.

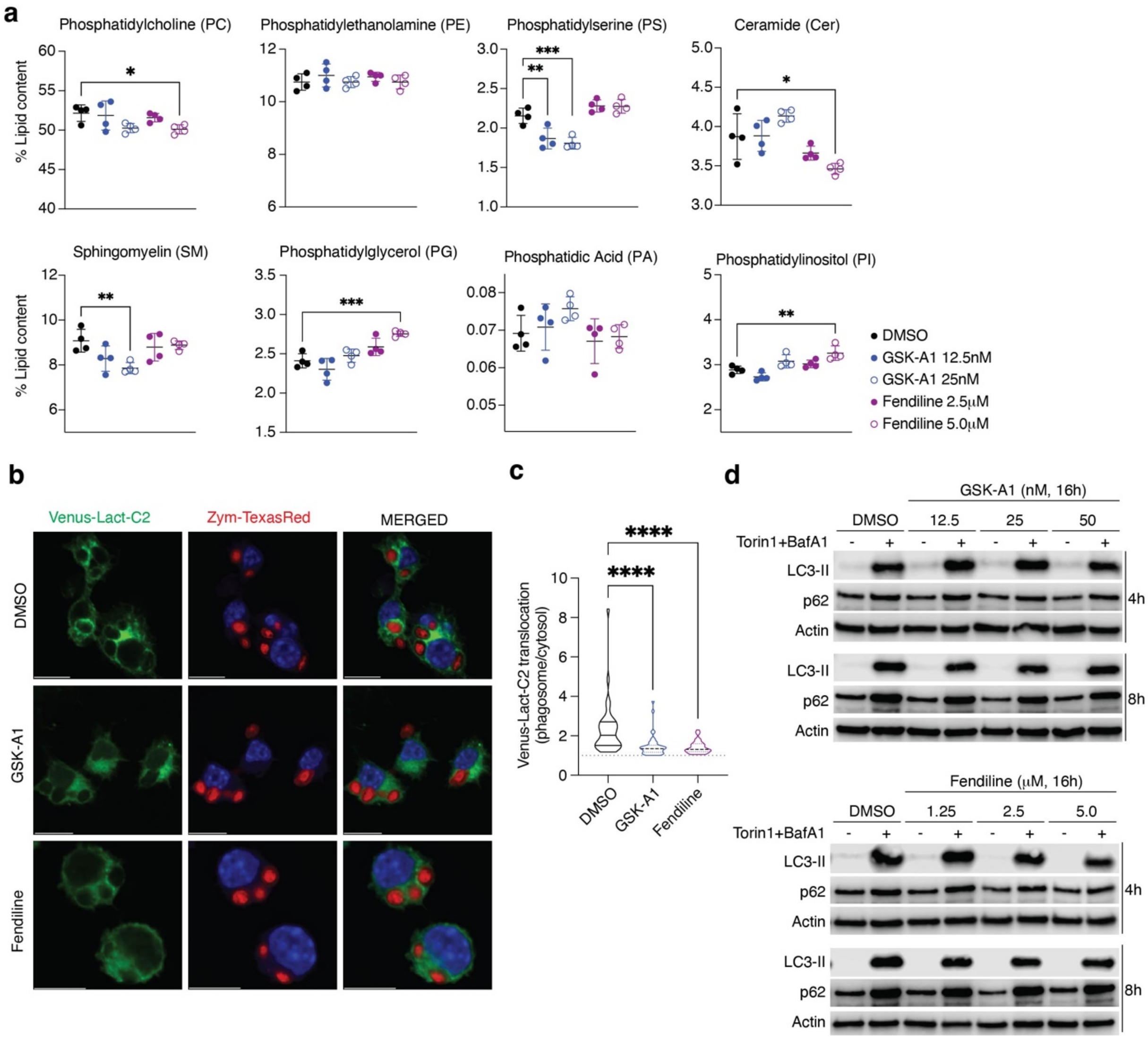


**Extended Data Figure 6: Effects of GSK-A1 and Fendiline as inhibitors of PS metabolism.** (**a**) RAW264.7 cells were pretreated as indicated (16h) and evaluated by lipidomics. Shown values were normalized to total lipid content per each sample. Data are means ± SD of 4 biological replicates in one experiment. (**b, c**) RAW264.7 stably expressing the PS-probe Venus-LACT-C2 pre-treated with GSK-A1 or Fendiline as indicated (16h) were fed Zymosan-TexasRed (Zym-TexasRed, 30 min). (b) Representative confocal images and (c) cumulative data of PS-probe enrichment in the phagosome membranes relative to cytosol in n>30 phagosomes in two independent experiments. (**d**) RAW264.7 cells were pre-treated with GSK-A1 or Fendiline for 16h at the indicated concentrations. Canonical autophagy was analyzed upon Torin1 (1μM) plus

BafilomycinA1 (20nM) treatment for 4h or 8h as indicated. Shown is representative immunoblot analysis of 3 independent experiments. * $P < 0.05$, ** $P < 0.01$, *** $P < 0.001$, **** $P < 0.0001$ by two-sided Student's t test.

Extended Data Figure 7. Boada-Romero et al.

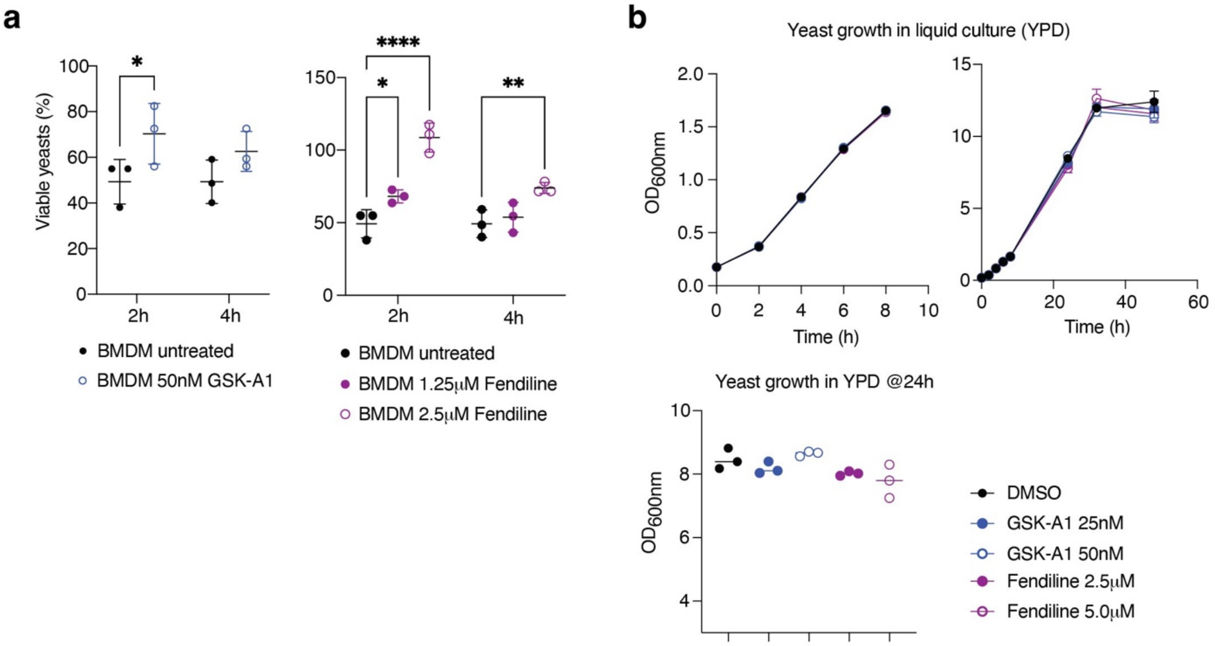


**Extended Data Figure 7: Fendiline blocks LAP**. (**a**) Yeast killing capacity after 2h and 4h of yeast engulfment by bone marrow derived macrophages (BMDM) pretreated with inhibitors as indicated (16h). Values normalized to yeast recovered after 1h of engulfment per cell line (time 0h). Untreated control values are shared in both graphs. (**b**) Growth curves ($OD_{600nm}$) of *Saccharomyces cerevisiae* at 30°C in liquid Yeast-Peptone-Dextrose (YPD) media supplemented with GSK-A1 or Fendiline at the indicated concentrations. Short-term, long-term and $OD_{600nm}$ values at 24h are shown. Data are means ± SD of three biological replicates in one representative of 3 independent experiment, n=3. * *P* <0.05, ** *P* <0.01, **** *P* <0.0001 by two-sided Student's t test.

Extended Data Figure 8. Boada-Romero et al.

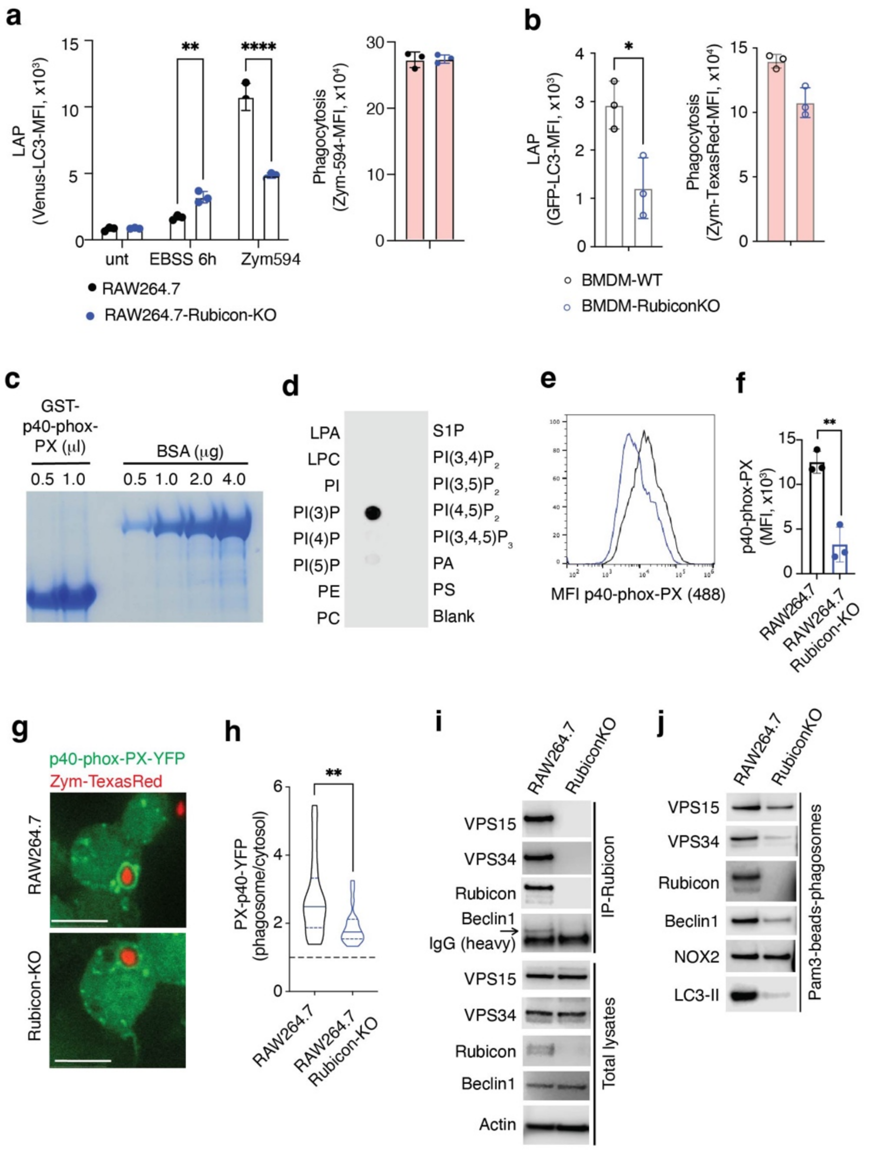


**Extended Data Figure 8: Rubicon mediates PI3P generation in the phagosomes**. (**a**) Flow cytometry analysis of retained Venus-LC3 in RAW264.7 or RAW264.7-Rubicon-KO cells stably expressing Venus-LC3. Cells were starved (Earle's Balanced Salt Solution, EBSS, 6h) or fed Zymosan-AlexaFluor594 particles (1h). Retained Venus-LC3 levels were analyzed after digitonin treatment. (**b**) Flow cytometry analysis of retained GFP-LC3 in BMDM cells from Rubicon-KO mice or littermate controls. Cells were fed Zymosan-TexasRed (1h) and treated as in (a). Right graphs in (a, b) show phagocytosis. (**c, d**) Recombinant GST-p40-phox-PX produced in bacteria was analyzed by (c) SDS-PAGE and Coomassie blue staining alongside bovine serum albumin

(BSA) standards or (d) lipid binding specificity using lipid strip. LPA: lysophosphatidic acid; LPC: lysophosphatidylcholine; PI: phosphatidylinositol; PE: phosphatidylethanolamine; PC: phosphatidylcholine; S1P: sphingosine-1-phosphate; PA: phosphatidic acid; PS: phosphatidylserine, n=2. (**e, f**) Flow cytometry analysis of PI3P in cells treated as in (a) and stained using recombinant GST-p40-phox-PX probe, anti-GST and Alexa-488 secondary antibodies. (e) PI3P levels as MFI-Alexa-488 histogram (e) or (f) mean of Alexa-488 in TexasRed$^+$ population. (**g, h**) RAW264.7 and RAW264.7-Rubicon-KO cells stably expressing the PI3P probe p40-phox-PX-Venus were fed Zymosan-TexasRed (30min). (g) Representative confocal images and (h) violin-plots of PI3P-probe enrichment at the phagosome membrane relative to cytosolic signal (n>10 phagosomes) in a representative experiment. (**i, j**) Immunoblot analysis of (i) anti-Rubicon immunoprecipitates (IP) of cell lysates and their corresponding total cell lysates and (j) lysates from isolated Pam3csk4-beads containing phagosomes from RAW264.7 or RAW264.7-Rubicon-KO cells. (a, b, f) Data are means ± SD of mean fluorescent intensity (MFI) in three biological replicates, representative of 3 independent experiments. * $P$ <0.05, ** $P$ <0.01, **** $P$ <0.0001 by two-sided Student's t test.

Extended Data Figure 9. Boada-Romero et al.

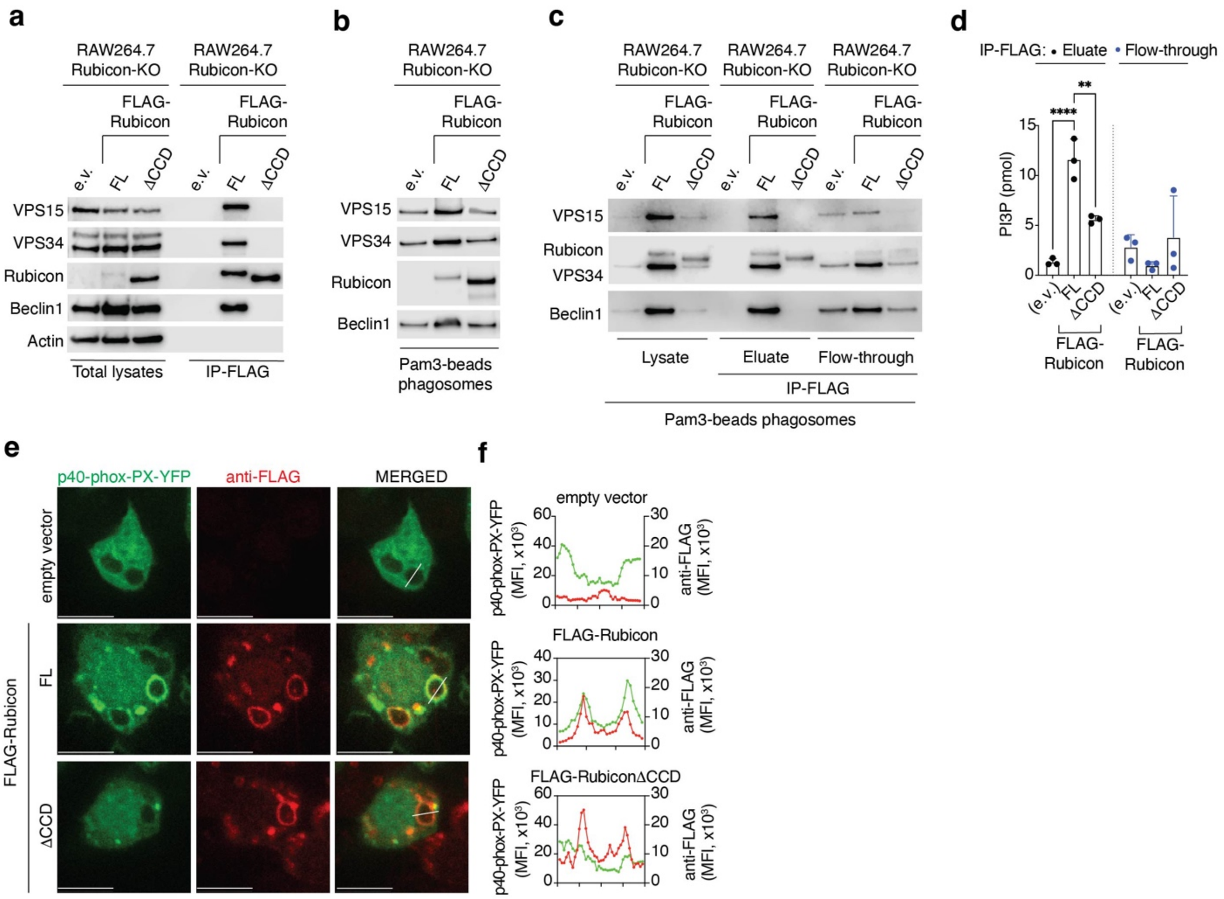


**Extended Data Figure 9: Rubicon-containing PI3KC3 complex holds lipid kinase activity on phagosomes**. (**a**-**c**) Immunoblot analysis of RAW264.7-Rubicon-KO cells stably expressing FLAG-tagged full-length Rubicon, ΔCCD mutant version (lacking the Beclin1-interacting region, amino acids 490-542), or empty vector (e.v.). (a) Total cellular lysates from cells were subjected to anti-FLAG immunoprecipitation (IP). (b, c) Lysates from isolated Pam3csk4-bead-containing phagosomes were subjected to anti-FLAG IPs; eluates and flow-through were run in SDS-PAGE. (**d**) Eluates and flow-through fractions from (c) were assayed for PI3P generation *in vitro*. (**e, f**) Cells as in (a) and stably expressing PI3P probe p40-phox-PX-Venus were fed Zymosan (30min) and stained for FLAG expression. (e) Representative confocal images, (f) mean fluorescence intensity (MFI) of p40-phox-PX-Venus (in green) and FLAG signal (in red) across the region of interest (white lines in images). ** *P* <0.01, **** *P* <0.0001 by two-sided Student's t test.

Extended Data Figure 10. Boada-Romero et al.

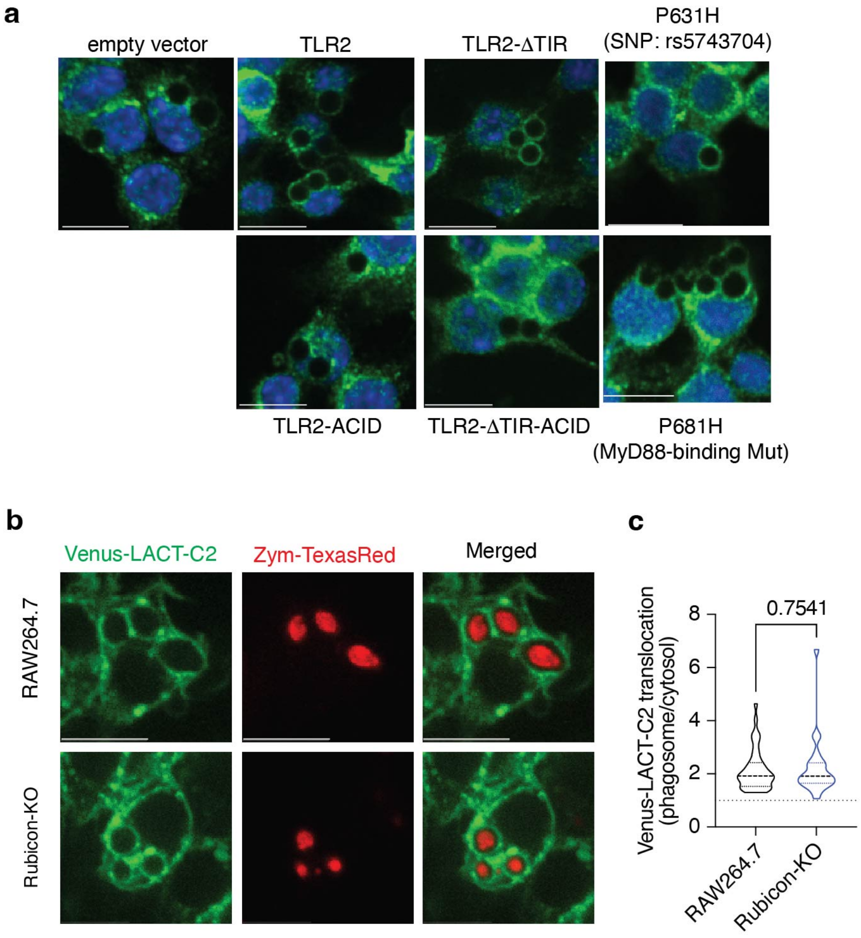


**Extended Data Figure 10: The TLR2-PS-Rubicon axis**. (**a**) RAW264.7-sg-TLR2-KO cells were transduced to express full length TLR2 or TLR2 lacking the TIR domain (TLR2ΔTIR), in either wild-type (WT), K628E-R629D-K630E-K632E-K633E (ACID), P631H (corresponding to human SNP rs5743704), or P681H mutant TLR2. Transduction with empty vector (e.v.) serves as a negative control. Cells were fed Pam3csk4-beads (30min) and Rubicon translocation was determined by immunofluorescence. (**b, c**) RAW264.7 and RAW264.7-Rubicon-KO cells stably expressing the PS-probe Venus-LACT-C2 were fed Zymosan-TexasRed (Zym-TexasRed, 30 min). (a, b) Representative confocal images and (c) violin-plots showing the enrichment of the PS-probe at the phagosome membrane relative to the cytosolic level (n>20 phagosomes). Violin plots display median and upper and lower quartiles. *P-value* by two-sided Student's t test is not significant.

Extended Data Figure 11. Boada-Romero et al.

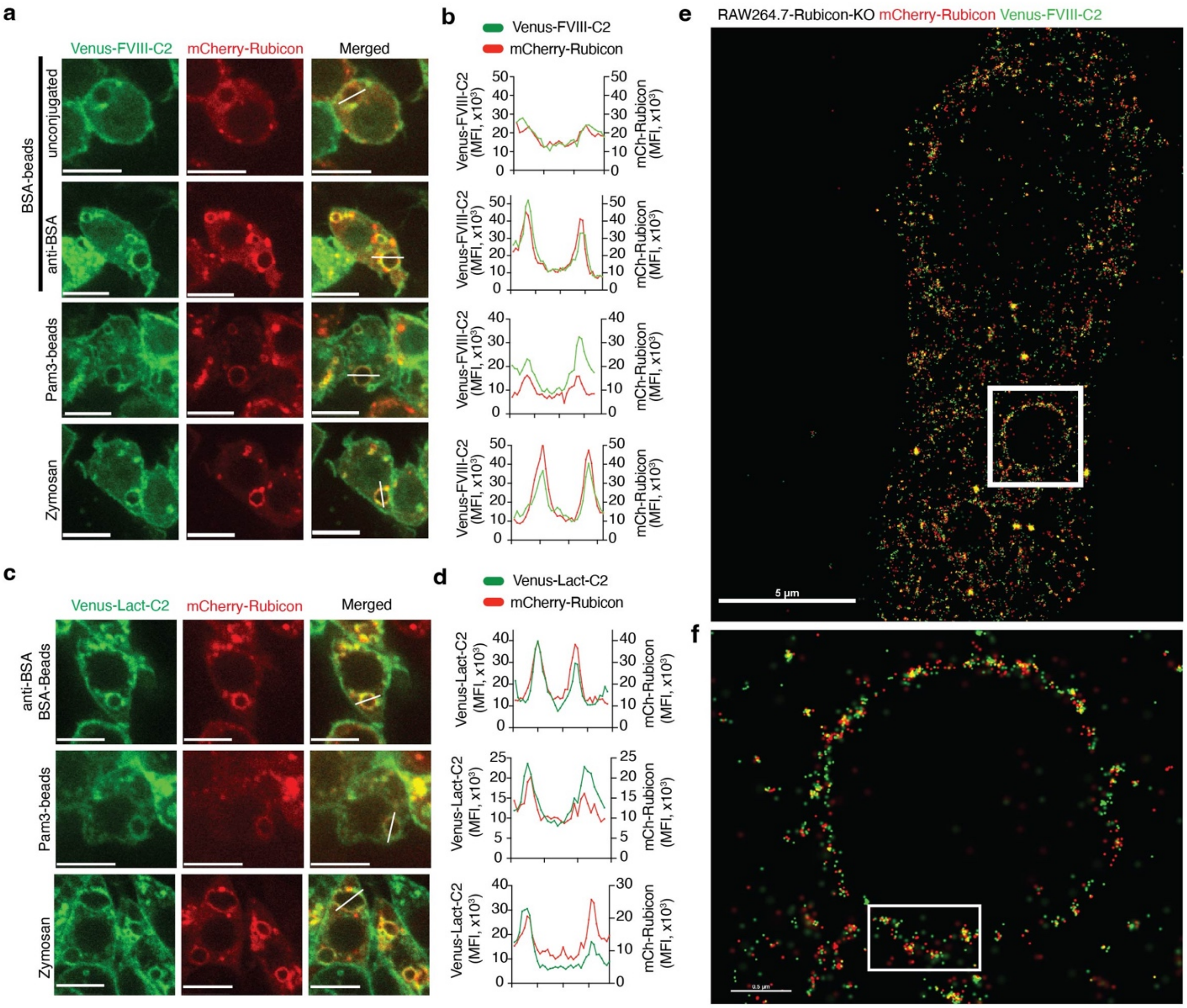


**Extended Data Figure 11: Colocalization of Rubicon and PS at the phagosome membrane**. RAW264.7-Rubicon-KO cells transduced for Rubicon expression (mCherry-Rubicon) and stably expressing fluorescent PS-probes Venus-FVIII-C2 (**a, b**) or Venus-Lact-C2 (**c, d**) were subjected to different phagocytic stimuli (30 min). Representative images (a, c) and profiles (b, d) of MFI of Venus and mCherry channels across the regions of interest (white lines in images) are shown. (**e, f**) Stochastic optical reconstruction microscopy (STORM) images of cells as in (a, b) fed Zymosan (15 min). Shown is a complete cell (e), where the boxed insert highlights a phagosome enlarged in (f). Insert in panel (f) is depicted in Fig. 5d as a sample of phagosome membrane.

Extended Data Figure 12. Boada-Romero et al.

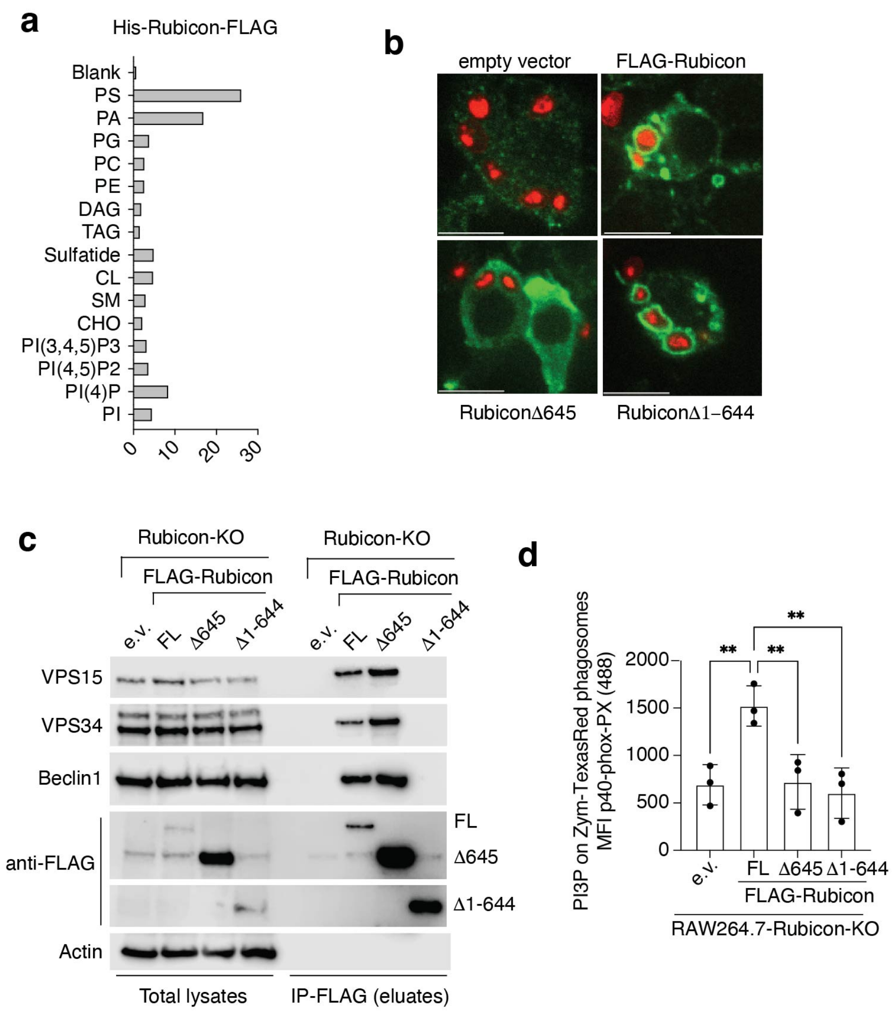


**Extended Data Figure 12: Translocation of Rubicon to phagosomes and stimulation of lipid kinase activity are independent events.** (**a**) Lipid strip showing the binding specificity of full-length Rubicon recombinantly produced in insect cells. Levels were quantified by densitometry and normalized to the *Blank* (dotting buffer). PS: phosphatidylserine; PA: phosphatidyl acid; PG: phosphatidylglycerol; PC: phosphatidylcholine; PE: phosphatidylethanolamine; DAG: diacyl-glycerol; TAG: triacyl-glycerol; CL: cardiolipin; SM: sphingomyelin; CHO: cholesterol; PI: phosphatidylinositol. (**b**, **c**) RAW264.7-Rubicon-KO cells stably expressing FLAG-tagged Rubicon, Δ645, Δ1-644, or empty vector (e.v.) were (b) fed Zymosan (30min) and imaged for FLAG staining or (c) lysed and total cell lysates were subjected to anti-FLAG immunoprecipitation (IP). Shown are representative images and immunoblots of 3 independent experiments. (**d**) Cells as in (b) were fed Zymosan-TexasRed particles, and after digitonin treatment, PI3P levels in Zymosan-TexasRed$^{+}$ cells were analyzed using recombinant GST-p40-phox-PX probe, anti-GST and Alexa488 secondary antibodies. Data are means ± SD of mean Alexa488 fluorescence in TexasRed$^{+}$ population of three biological replicates. Representative of 3 independent experiments. ** $P$ <0.01 by two-sided Student's t test.

Extended Data Figure 13. Boada-Romero et al.

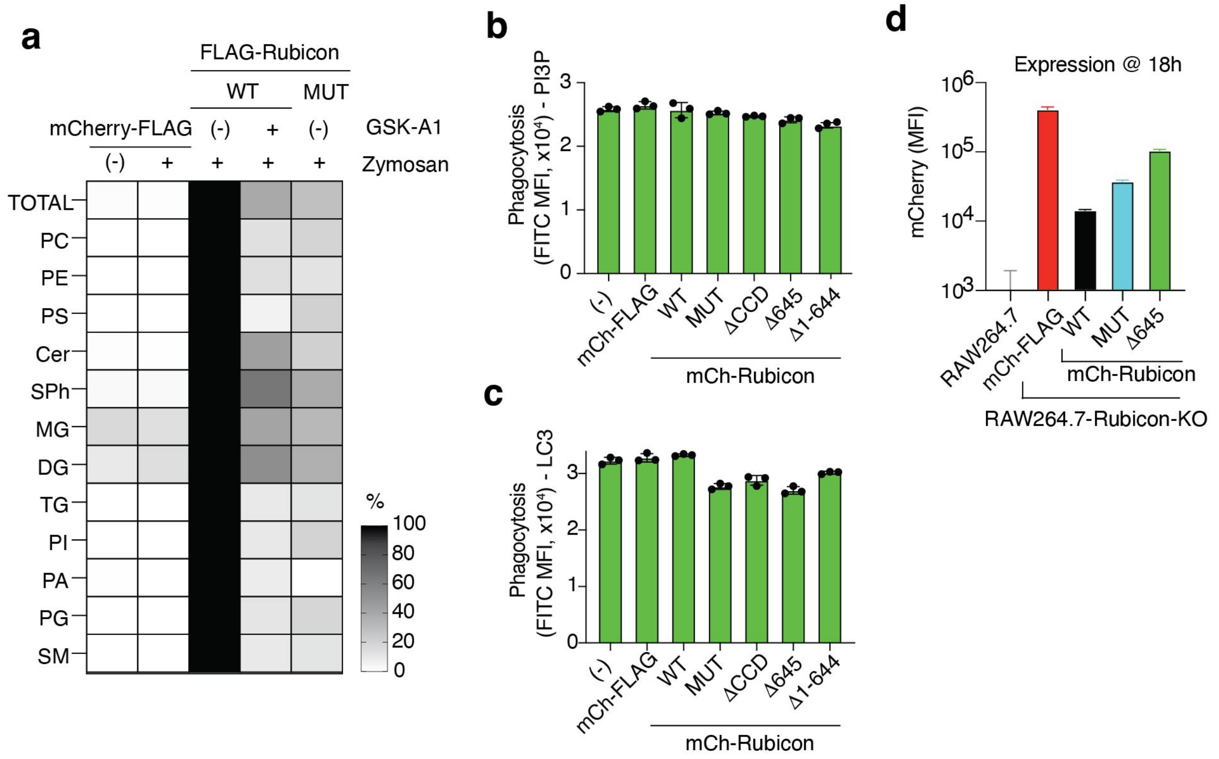


**Extended Data Figure 13: Rubicon-K718E-R719E-R721E is not able to bind to lipids.** (**a**) RAW264.7-Rubicon-KO cells stably expressing FLAG-Rubicon in WT or K718E-R719E-R721E (MUT) version were pretreated with GSK-A1 (25nM, 16h) and then fed Zymosan particles as indicated. Cells were mechanically lysed, subjected to anti-FLAG immunoprecipitation (IP) followed by lipidomics. Total lipid content and several lipid classes are represented, values normalized to WT-Zymosan. PC: phosphatidylcholine; PE: phosphatidylethanolamine; PS: phosphatidylserine; Cer: ceramide; SPh: sphingosine; MG: monoacyl-glycerol; DG: diacyl-glycerol; TG: triacyl-glycerol; PI: phosphatidylinositol; PA: phosphatidyl acid; PG: phosphatidylglycerol; SM: sphingomyelin. (**b, c**) Phagocytosis of Zymosan-FITC particles (as mean fluorescent intensity, MFI, in $FITC^+$ cells) in indicated cells from experiment in Fig. 6e and 6f. (**d**) mCherry expression (as MFI values) in acidification experiments (at 18h) in Fig 6g. Data are means ± SD of 3 (b, c) or eight (d) biological replicates. Each representative of 3 independent experiments. mCherry-FLAG (mCh-FLAG) is used as negative control.

Extended Data Figure 14. Boada-Romero et al.

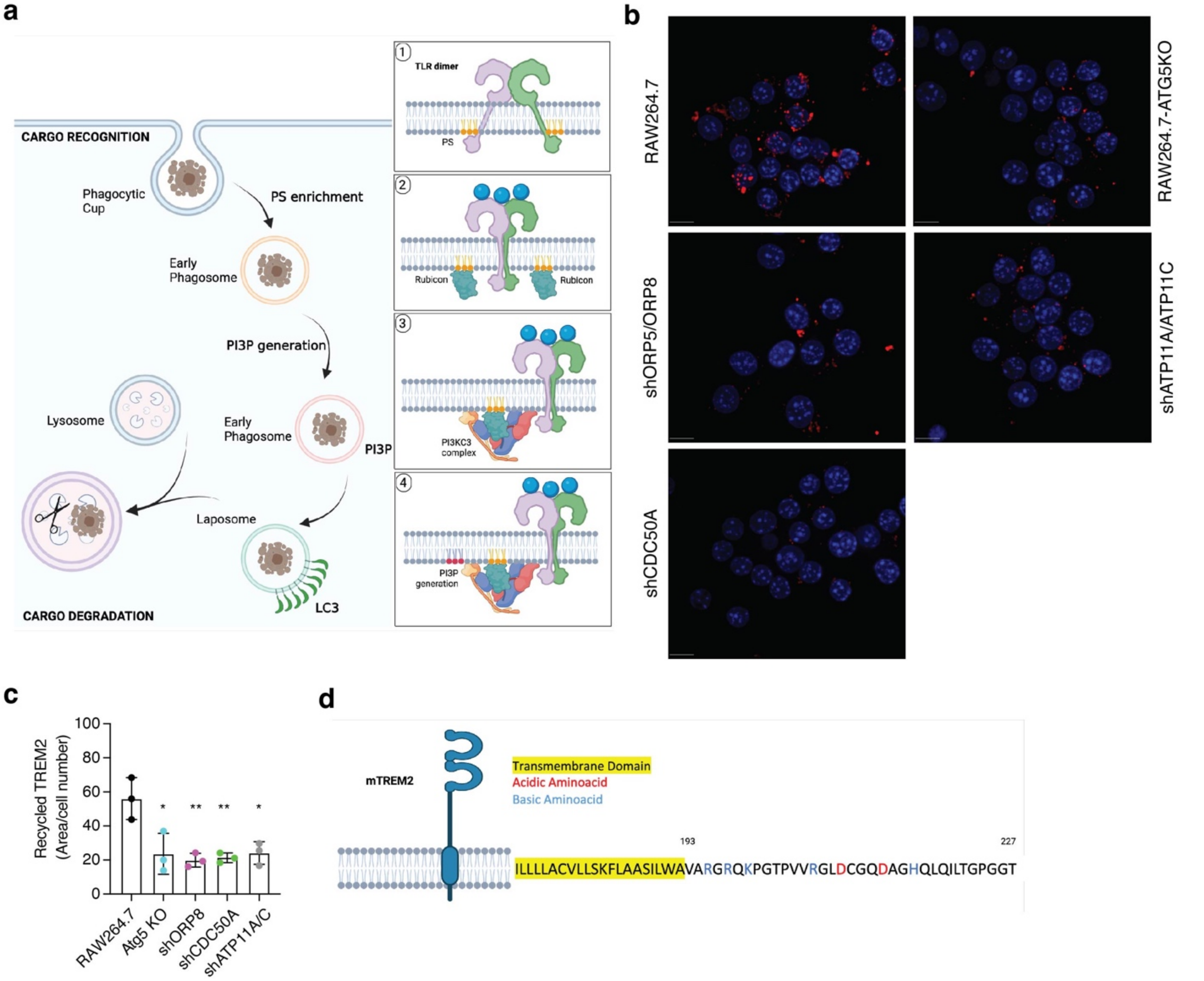


**Extended Data Figure 14: The Role of PS in LAP and LANDO.** (**a**) Model of LC3-associated phagocytosis. Overview (left) and proposed detailed mechanism (right). Sensing of cargo promotes the (1) enrichment of phagocytic receptors at the phagocytic cup, and phosphatidylserine (PS) clustered by receptor cytosolic domains leads to an enrichment of PS in the early phagosome. (2) PS attracts and docks Rubicon to the phagosome membrane that (3) favors the assembly of Rubicon-containing PI3KC3 complexes in the early phagosome, (4) stimulating the generation of PI3P in the early phagosome. Presence of PI3P recruits the LC3 lipidation machinery, assisting fusion of the phagosome to lysosomes to facilitate cargo degradation. (**b, c**) RAW264.7-Venus-LC3 cells silenced for ORP5 and ORP8, CDC50A, or ATP11A and ATP11C using stably transduced shRNAs (shORP5/8, shCDC50A, and shATP11A/C) were assessed for LANDO-dependent TREM2 receptor recycling. RAW264.7-Atg5-KO cells benchmark LANDO deficiency. (b) Representative pictures and (c) quantification of recycled TREM2 per cell number in 3 fields in one representative experiment (n=3). * $P$ <0.05, ** $P$ <0.01 by two-sided Student's t test. (d) Scheme of TREM2 (mouse, Uniprot: Q99NH8) depicted at the plasma membrane. Transmembrane domains are highlighted in yellow; acidic and basic amino acid are red and blue, respectively. Numbers indicate amino acid position.

Extended Data Table 1. Peptide Synthesized for lipid bilayer experiments.

| Name | Uniprot, ID region | Wild-type, Mutations | Sequence |
|---|---|---|---|
| HisX6-mouseCD16tail-PEG-Biotin | P08508, 236-261 | Wild-type | HHHHHHRRNLQTPREYWRKSLSIRKHQAPQDK-PEG-Biotin |
| HisX6-mouseCD16tail-RR-PEG-Biotin | P08508, 236-261 | R236E, R237E | HHHHHHEENLQTPREYWRKSLSIRKHQAPQDK-PEG-Biotin |
| HisX6-mouseCD16tail-RK-PEG-Biotin | P08508, 236-261 | R247E, K248D | HHHHHHRRNLQTPREYWEDSLSIRKHQAPQDK-PEG-Biotin |
| HisX6-mouseCD16tail-RKH-PEG-Biotin | P08508, 236-261 | R253E, K254D, H255D | HHHHHHRRNLQTPREYWRKSLSIEDDQAPQDK-PEG-Biotin |
| HisX6-mouseCD16tail-ALL-PEG-Biotin | P08508, 236-261 | R236E, R237E, R247E, K248D, R253E, K254D, H255D | HHHHHHEENLQTPREYWEDSLSIEDDQAPQDK-PEG-Biotin |
| HisX6-humanCD16-PEG-Biotin | P08637, 230-254 | Wild-type | HHHHHHKTNIRSSTRDWKDHKFKWRKDPQDK-PEG-Biotin |
| HisX6-humanCD16-ALL-PEG-Biotin | P08637, 230-254 | K230D, R234E, K241D, H243D, K244D, K246D, R248E, K249D | HHHHHHDTNIESSTRDWDDDDFDWEDDPQDK-PEG-Biotin |
| HisX6-humanTIM4tail-PEG-Biotin | Q96H15, 336-378 | Wild-type | HHHHHHLRGKLMETYCSQKHTRLDYIGDSKNVLNDVQHGREDEDGLFTL-PEG-Biotin |
| HisX6-humanTIM4tail-ALL PEG-Biotin | Q96H15, 336-378 | R337E, K339D, K348D, K349D, R351E, H367D, R369E | HHHHHHLEGDLMETYCSQDDTELDYIGDSKNVLNDVQDGEEDEDGLFTL-PEG-Biotin |
