## Supplementary Table1 for "Phosphatidylserine clustering by membrane receptors triggers LC3-associated phagocytosis"

| PLASMID | BACKBONE | SOURCE | PRIMERS |
|---|---|---|---|
| *Entry vectors, controls, viral helpers* | | | |
| pMXs-empty_vector-BLAST (entry plasmid) | pMXs-_-IRES-BLAST | Cell Biolabs, Inc; RTV-106 | N.A. |
| pMXs-FLAG-mCherry (control plasmid) | pMXs-_-IRES-BLAST | This study | FLAG: fw - gggcccggatccaccatgggcgactacaaagacgatgacgacaagtccggaggctccgg;<br>FLAG: rev - gggcccacgcgtcgccgccggagcctccggagcctccggacttgtcgtcatcgtctttg.<br>mCherry: fw - gggcccacgcgtccggcggaatgggcgtgagcaagggcgaggaggataac;<br>mCherry: rev - gggcccgcggccgctttacttgtacagctcgtccatgc. |
| pCL-AMPHO | | Imgenex, 10046P | N.A. |
| VSV-G | | Addgene#8454 | N.A. |
| PAX2 | | Addgene#12260 | N.A. |
| *Reporters: LAP, PS probe and PI3P probe* | | | |
| pMXs-Venus-LC3-IRES-BLAST (autophagy pathway reporter) | pMXs-_-IRES-BLAST | This study<br><br>Source<br>Addgene#21074 | Venus: fw - gggcccggatccaccatgggcagcaagggcgaggagctgttc;<br>Venus: rev - gggccctcatgacgccgccggacttgtacagctcgtccatgc.<br>ratLC3: fw - gggccctcatgagcggcggcatgccgtccgagaagacc;<br>ratLC3: rev - gggcccgcggccgctttacacagccagtgctgtcccg. |
| pMXs-Venus-LACT-C2 (PS reporter) | pMXs-_-IRES-BLAST | This study<br><br>Lact cDNA from mouse cDNA library | Venus N-term: fw - gggcccggatccaccatgggcagcaagggcgaggagctgttc;<br>Venus N-term: rev - gggcccacgcgtcgccgccggacttgtacagctcgtccatgc.<br>LACT-C2: fw - gggccacgcgtccggcggaggatgttctgagcccctgggcctgaag;<br>LACT-C2: rev - gggcccgcggccgctttaacagcccagcagctccaggcgcaggg. |
| pMXs-Venus-FVIII-C2 (PS reporter) | pMXs-_-IRES-BLAST | This study, FVIII cDNA from mouse cDNA library | fw - gggccacgcgtccggcggaagttgcagcataccattgggaatggaaag;<br>rev - gggcccgcggccgctttagtattgctgctgggcctcacatcctagaatctc |
| pMXs-p40phox-Venus (PI3P reporter) | pMXs-_-IRES-BLAST | This study; Ncf4 cDNA from mouse cDNA library | PX: fw – gggcccagatctaccatgggcctggcccagcagctgcgatcag;<br>PX: rev – gggcccgcggccgctttagcggagtgcctggggcacctgctc.<br>Venus C-term: fw - GGGCCacgcgtccGGCGGAATGGGCAGCAAGGGCGAGGAGCTGTTC;<br>Venus C-term: rev - gggcccgcggccgcTTTA CTTGTACAGCTCGTCCATGC. |
| *Rubicon Constructs* | | | |
| pMXs-FLAG-Rubicon;<br>pMXs-mCherry-Rubicon | pMXs-_-IRES-BLAST | This study; Rubicon cDNA from Addgene#21636 | fw - gggccacgcgtccggcggaatgcgtccggagggcgcggg;<br>rev - gggcccgcggccgctttaggtggtctctaggacggtgg. |
| pMXs-FLAG-RubiconDCCD;<br>pMXs-mCherry-RubiconDCCD | pMXs-_-IRES-BLAST | This study | N-terminal: fw - gggccacgcgtccggcggaatgcgtccggagggcgcggg;<br>N-terminal: rev - gggccccaattgggcaatcaaggactcggagatgctgaagtgg;<br>C-terminal: fw - gggccccaattgactgagaatggaagcttccgggtcacc;<br>C-terminal: rev - gggcccgcggccgctttaggtggtctctaggacggtgg. |
| pMXs-mCherry-RubiconD645 | pMXs-_-IRES-BLAST | This study | fw - gggccacgcgtccggcggaatgcgtccggagggcgcggg;<br>rev - gggcccgcggccgctttacttctgtggagcatcatgctctgggactagcc. |
| pMXs-mCherry-RubiconD1-644 | pMXs-_-IRES-BLAST | This study | fw - gggcccacgcgtccggcggactcctacccatccccgactccctgccc;<br>rev - gggcccgcggccgctttaggtggtctctaggacggtgg. |
| pMXs-FLAG-Rubicon-K718E-R719E-R721E;<br>pMXs-mCherry-Rubicon-K718E-R719E-R721E | pMXs-_-IRES-BLAST | This study | top - ccggacggaccccgactatatcGAGGAActgGAGtactgtgagtacctaggcaag;<br>bottom - cttgcctaggtactcacagtaCTCcagTTCCTCgatatagtcggggtccgtccgg. |
| *TLR2 constructs* | | | |
| pMXs-TLR2-sgi | pMXs-_-IRES-BLAST | This study; Tlr2 cDNA from Addgene#13083; | TLR2: fw – gggcccatgcataccatgggccgagctctttggctcttctggatcttgg;<br>TLR2: rev – gggcccgcggccgctttaggactttattgcagttctcagatttaccc.<br>sgi: top – ggtgtgtgatggccgctccCGAtctttcacctctattccc;<br>sgi: bottom – gggaatagaggtgaaagaTCGggagcggccatcacacacc. |
| pMXs-TLR2-sgi-K628E-R629D-K630E-K632E-K633E | pMXs-_-IRES-BLAST | This study | top – atgtgggcgtggctccaggccGagGACGagcccGagGaagctccctgcagggacgtttgc;<br>bottom – gcaaacgtccctgcagggagcttCctCgggctCGTCctCggcctggagccacgcccacat |
| pMXs-TLR2-sgi-K628A-R629A-K630A-K632A-K633A | pMXs-_-IRES-BLAST | This study | top – atgtgggcgtggctccaggccGCCGCTGCCcccGCGGCAgctccctgcagggacgtttgc;<br>bottom – gcaaacgtccctgcagggagcTGCCGCgggGGCAGCGGCggcctggagccacgcccacat |
| pMXs-TLR2-sgi-P631H | pMXs-_-IRES-BLAST | This study | top - ggctccaggccaagaggaagcAcaagaaagctccctgcaggg;<br>bottom – ccctgcagggagctttcttgTgcttcctcttggcctggagcc |
| pMXs-TLR2-sgi-P681H | pMXs-_-IRES-BLAST | This study | top – ctccacaagcgggacttcgttcACggcaaatggatcattgacaac;<br>bottom – gttgtcaatgatccatttgccGTgaacgaagtcccgcttgtggag; |
| pMXs-TLR2-sgi-DTIR (D640) | pMXs-_-IRES-BLAST | This study | fw – gggcccatgcataccatgggccgagctctttggctcttctggatcttgg;<br>rev – gggcccgcggccgctttaaacgtccctgcagggagctttcttgggc |
| pMXs-TLR2-sgi-DTIR-K628E-R629D-K630E-K632E-K633E | pMXs-_-IRES-BLAST | This study | fw – gggcccatgcataccatgggccgagctctttggctcttctggatcttgg;<br>rev – gggcccgcggccgctttaaacgtccctgcagggagcttcctcgggc |
| *Recombinant Expression* | | | |
| pGEX-4T1 (empty vector, for GST only) | pGEX-4T1 | Cytiva 28-9545-49 | N.A. |
| pGEX-4T1-p40-phox | pGEX-4T1 | This study | fw – gggcccagatctaccatgggcctggcccagcagctgcgatcag;<br>rev – gggcccgcggccgctttagcggagtgcctggggcacctgctc. |
| pGEX-4T1-FLAG-Rubicon-D1-644 | pGEX-4T1 | This study | fw - gggcccggatccaccatgggcgactacaaagacgatgacgacaagtccggaggctccgg;<br>rev - gggcccgcggccgctttaggtggtctctaggacggtgg. |
| pGEX-4T1-TLR2-ID(589-784);<br>pGEX-4T1-TLR2-ID-K628E-R629D-K630E-K632E-K633E;<br>pGEX-4T1-TLR2-ID-K628A-R629A-K630A-K632A-K633A | pGEX-4T1 | This study, pMXs as cDNA source | fw – ggccggatcctccggaggctccggaggctccggcgggtgccaccatttccacggactgtg;<br>rev – gggcccgcggccgctttaggactttattgcagttctcagatttaccc |
| pFAST-BAC-His6X-Rubicon-FLAG | pFAST-Bac-HT | This study | Fw - gggcccggatccaccATGcgtccggagggcgcgggaatggatc;<br>rev -; gggcccacgcgtcGCCGCCggtggtctctaggacggtggcctc<br>FLAG (Cterm): wt - GGGCCCacgcgtccGGAGGCTCCGGAGGCtccggcggaGACTACAAAGACGATGACGAC<br>FLAG (Cterm) rev - gggcccgcggccgcTTTACTTGTCGTCATCGTCTTTGTAGTCtccgccggaGCCTCCGG |
| pNIC-His6X-TLR2-ID-FLAG;<br>pNIC-His6X-TLR2-ID-K628E-R629D-K630E-K632E-K633E-FLAG | pNIX-Bsa4 | This study | fw - gggccccatatgcatcatcaccatcaccactgccaccatttccacggactgtggtacc;<br>rev - cccggggcggccgctttacttgtcgtcatcgtctttgtagtcgcctccggactttattgcagttctcagatttaccc. |
| *shRNA for Knockdown* | | | |
| pLKO-sh-mouse-OSBPL5/ORP5 | pLKO.1 | SIGMA | TRCN0000105111 |
| pLKO-sh-mouse-OSBPL8/RP8 | pLKO.1 | SIGMA | TRCN0000105248 |
| pLKO-sh-mouse-ATP11A | pLKO.1 | SIGMA | TRCN0000101533 |
| pLKO-sh-mouse-ATP11C | pLKO.1 | SIGMA | TRCN0000101851 |
| pLKO-sh-mouse-TMEM30A/CDC50A | pLKO.1 | SIGMA | TRCN0000317704 |
| *CRISPR-Cas9 for Knock-out* | | | |
| LentiCRISPR-v2-empty vector | LentiV2 | Addgene#98290 | N.A. |
| LentiCRISPR-v2-sgTLR2 | LentiV2 | This study | top – caccgggtgtgtgatggccgctcc;<br>bottom - aaacggagcggccatcacacaccc |
